# Highly active chromosome regions preferentially associate with two perispeckle networks that partition the interchromatin space

**DOI:** 10.1101/2025.08.23.671945

**Authors:** Neha Chivukula Venkata, Jiah Kim, Gabriel Faber, Saireet Misra, Atabek Bektash, Pankaj Chaturvedi, Gabriela Hernanadez, Joseph Dopie, Masato T. Kanemaki, Kyu Young Han, Yaron Shav Tal, Andrew S. Belmont

**Affiliations:** Department of Cell and Developmental Biology, University of Illinois at Urbana-Champaign, Urbana, IL; The Mina and Everard Goodman Faculty of Life Sciences, Bar-Ilan University, Ramat Gan 5290002, Israel; National Institute of Genetics, Research Organization of Information and Systems (ROIS), Mishima, Shizuoka 411-8540, Japan; Graduate Institute for Advanced Studies, SOKENDAI, Mishima, Shizuoka 411-8540, Japan; Department of Biological Science, Graduate School of Science, The University of Tokyo, Tokyo 113-0033, Japan; CREOL, The College of Optics and Photonics, University of Central Florida, Orlando, FL, USA; Center for Biophysics and Quantitative Biology, University of Illinois at Urbana-Champaign, Urbana, IL; Carl R. Woese Institute for Genomic Biology, University of Illinois at Urbana-Champaign, Urbana, IL

**Keywords:** Genome organization, active chromatin, interchromatin compartment, nuclear speckles

## Abstract

A subset of highly active chromosomal “hot zones” reproducibly positions adjacent to nuclear speckles (NS). Genes within these regions amplify their expression only with NS contact. However, gene expression differences inversely correlate with differences in NS distance, genome-wide. We hypothesized the existence of additional gene expression “niches” away from, but spatially correlated with, NS. Here we report the identification of two dynamic perispeckle patterns of protein concentrations extending outwards from NS and persisting even after NS are eliminated. Highly active chromosome regions which weakly associate with NS instead show close, NS-independent association with these perispeckle patterns. Additionally, transcripts from model intron-containing versus intronless genes associate differentially with these two patterns. While genes within NS-associated genomic regions are predominantly downregulated upon NS depletion, genes associated with perispeckle patterns are biased towards upregulation. We suggest the interchromatin space is partitioned into additional gene expression “niches”- surrounding and extending from NS - that may be involved in mRNA and gene dynamics.

## Introduction

Nuclear speckles (NS) were first suggested to act as a gene expression hub for a subset of active genes based on a survey of ∼20 active genes using fluorescence in situ hybridization (FISH) (Smith et al., 1999; Hall et al., 2006). Over the past several years, this model has now been validated by multiple orthogonal genome-wide approaches, including TSA-seq, SPRITE, and highly multiplexed immuno-FISH (Quinodoz et al., 2018; Chen et al., 2018; Su et al., 2020; Takei et al., 2021; Zhang et al., 2021; Gholamalamdari et al., 2025). A distinct subset of chromosomal regions-GC-rich, gene-dense, and enriched with shorter genes with fewer introns and house-keeping genes-show high frequency association with NS and mean NS distances of just several hundred nm (Chen et al., 2018). While several developmentally regulated genes have been shown to move from the nuclear lamina to NS after their gene activation, unexpectedly most of these Speckle-Associated Domains (SPADs) are highly conserved across multiple cell types (Zhang et al., 2021; Takei et al., 2021, 2023; Bhat et al., 2024). The most highly expressed genes show a pronounced enrichment in SPADs, as do CTCF binding sites, super-enhancers, and active marks such as H3K27ac, H3K9ac, and K3K79me3 (Chen et al., 2018; Yu et al., 2023b; Wang et al., 2021; Joo et al., 2023). In addition to this correlation of high levels of gene expression with NS association, live-cell imaging revealed a striking increase in *HSPA1* gene expression occurring within several minutes of gene movement and contact with NS (Khanna et al., 2014; Kim et al., 2020). Similar “amplification of gene expression” with NS association was inferred for several genes flanking the *HSPA1* locus (Kim et al., 2020), for the *HSPH1* gene (Zhang et al., 2021), and for several hundred p53-responsive genes (Alexander et al., 2021). Additionally, NS proximity has been shown to increase the efficiency of RNA splicing (Bhat et al., 2024).

While these results firmly establish NS as a nuclear hub for a subset of active genes, they raise the obvious next question of whether other nuclear compartments might similarly act as a hub for different subsets of active genes. Several previous observations led us to speculate that there might be additional nuclear compartments which act as local niches for gene expression.

First, we observed two types of chromosome regions with similarly elevated levels of gene expression; both types of these gene expression “hot-zones”, on average several hundred kbp in size, aligned with SON TSA-seq local maxima (Chen et al., 2018) (Gholamalamdari et al., 2025). Whereas high-amplitude, “Type I” SON TSA-seq peaks corresponded to chromosome regions strongly associated with NS, low-amplitude, “Type II” SON TSA-seq peaks corresponded to chromosome regions with weak association with NS (Gholamalamdari et al., 2025). Second, based on the shapes of the Type I SON peaks, aligned with corresponding Lamin B TSA-seq valleys, we inferred chromosome trajectories extending from the nuclear lamina and terminating adjacent to NS for the several hundred to thousand kbp lying between Type I SON TSA-seq peaks and the nearest LAD, as validated for several trajectories by DNA FISH (Chen et al., 2018). By analogy, we hypothesized similar chromosome trajectories extending from the nuclear lamina towards an unknown nuclear body/compartment that Type II SON TSA-seq peaks would associate with instead of NS. Third, chromosome regions shifting in different cell types from far to moderate distances to NS showed similar biases towards increases in gene expression as other chromosome regions that shifted from moderate to near distances to NS (Zhang et al., 2021). Fourth, live-cell imaging revealed repeated nucleation and then movement of small NS towards larger NS within DNA-depleted channels connecting the NS (Kim et al., 2019). Despite the known fast “constrained diffusion” of chromosome loci over radii of ∼500-1000 nm (Marshall et al., 1997; Chubb et al., 2002; Levi et al., 2005), these channels remained DNA-depleted over tens of minutes suggesting that some unknown non-chromatin component(s) might be occupying these channels.

Our observations showing an inverse relationship between changes in gene expression and changes in NS proximity across a wide range of NS distances created an apparent paradox when compared to our live-cell imaging observations which instead showed amplification of gene expression of certain genes only after actual microscopic contact with NS. To resolve this paradox, we hypothesized that just as certain genes might need to actually contact NS to enhance their gene expression, other genes might need to contact a different nuclear body/compartment to enhance their gene expression. If these putative additional nuclear bodies/compartments were themselves spatially correlated with NS, then a gene movement leading to actual contact with these nuclear bodies/compartments would appear by SON TSA-seq as a relative movement towards but not close to NS, resolving the apparent paradox.

Here we describe identification of several markers for each of two interwoven perispeckle staining patterns. Together, these perispeckle patterns meet our predictions of additional novel compartment(s) surrounding and extending from NS and serving as alternative gene expression hubs for a subset of highly active genes not associated with NS. The differential association of intron-rich versus intron-poor mRNAs with these two perispeckle staining patterns, together with the known roles of many of their marker proteins in RNA processing and/or RNA export, additionally suggests that these perispeckle staining patterns may define nuclear compartments that help spatially integrate the processes of transcription, RNA processing, and nuclear export.

## RESULTS

### Screening for potential markers of alternative “gene expression hubs”

As summarized in the Introduction, we postulated the existence of a nuclear compartment(s), itself spatially correlated with NS, that might act as an alternative hub for the several hundred kbp chromosome gene expression “hot-zones” that were not closely associated with NS (Fig. 1A). We imagined that such a compartment(s) might lie adjacent to and/or extend outwards from NS. Thus, shifts of chromosome regions towards this compartment(s) would then score as an increase in the NS TSA-seq score, which probes the proximity of chromosome regions to NS on an approximately micron-length scale.

**Figure 1:**
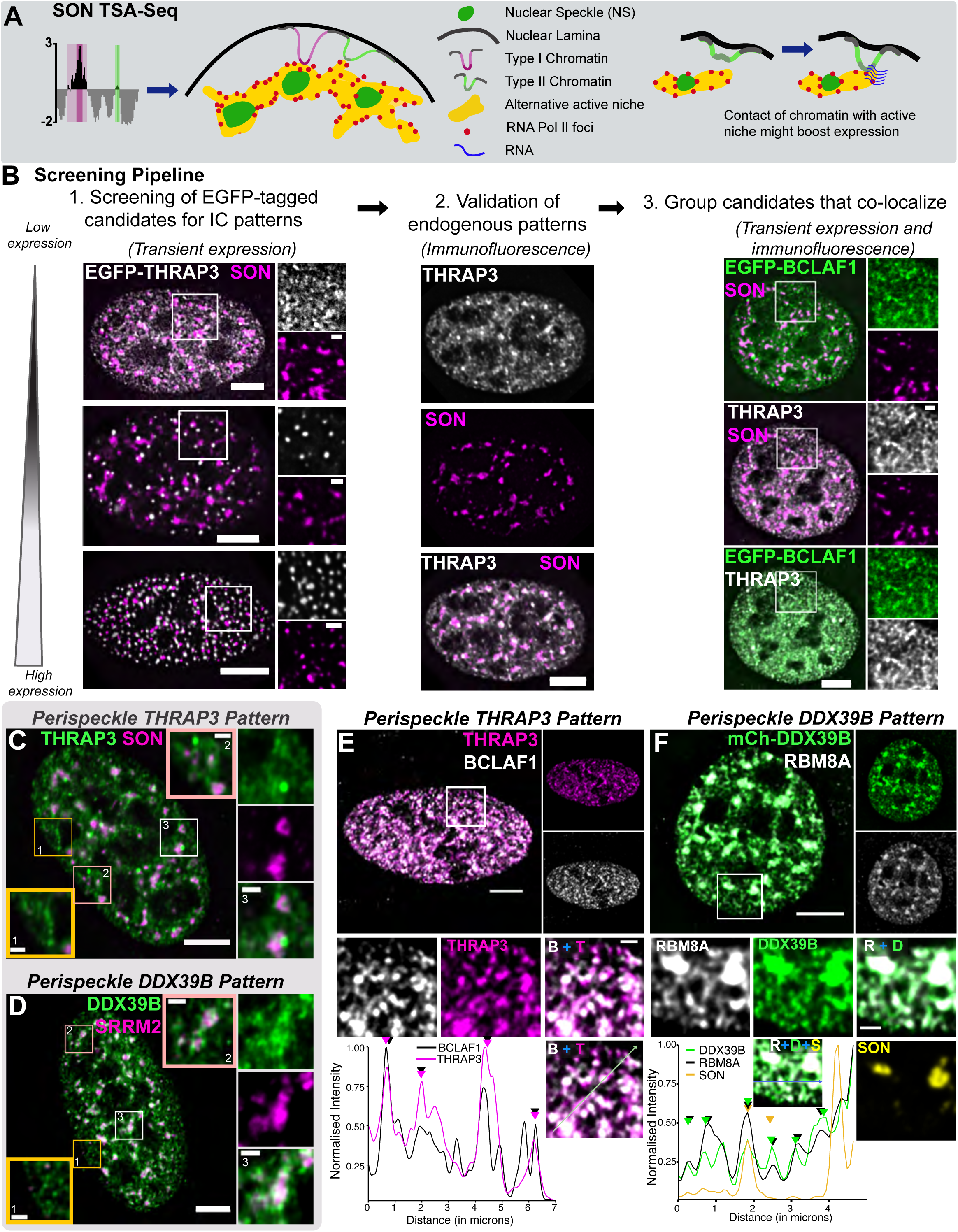
Screening for potential markers of alternative “gene expression hubs”. A. TSA-seq SON Type 1 (dark magenta) and Type 2 (dark green) peaks differ in their association with and average distance from NS. Type 2 peaks represent active genomic regions, weakly associated with NS, that might instead preferentially interact with an alternative nuclear active niche (yellow) present in the interchromatin (IC) space. Contact with this alternative active niche might boost gene expression (right panel). B. Screening pipeline for the identification of proteins that occupy IC space, illustrated with THRAP3 as an example candidate protein. (Left) Transient transfection of EGFP-tagged constructs (gray) in some cases showed varied distributions based on expression levels (gradient, left), which often accompanied changes in NS morphology. (Middle) Candidates showing a network-like pattern at low expression levels were validated by antibody staining of the endogenous protein. (Right) Candidate proteins were grouped into the same staining pattern if they showed extensive colocalization throughout the nucleus. C-D: Perispeckle THRAP3 Pattern (PSTP) (C) and Perispeckle DDX39B Pattern (PSDP) (D) represent two networks within the IC space that show protein concentrations in the vicinity of NS but also extend outwards in a network-type pattern. Insets: Box 1 (yellow borders): string of foci form linear thread-like structures; Box 2 (white borders): thread-like extensions at the nuclear periphery; Box 3 (pink borders): thread-like extensions connecting nearby NS. PSTP proteins are present at similar concentration within and outside NS, but PSDP proteins are concentrated inside NS as compared to the network outside of NS. E-F: Colocalization of PSTP (E) and PSDP (F) proteins. (E) Immunostaining of BCLAF1 (gray) relative to THRAP3 (magenta) and EGFP-SON (yellow). (F) Immunostaining of RBM8A (gray) relative to mCherry-DDX39B (green) and EGFP-SON (yellow). Line profiles along the arrows (shown in merged insets) show overlapping peaks (arrowheads) of BCLAF1 and THRAP3 (left) or RBM8A and DDX39B (right). Scalebars: 5 μm (main images) and 1 μm (insets).

We previously applied Tyramide Signal Amplification coupled to mass spectrometry (TSA-MS) to identify NS-enriched proteins (Dopie et al., 2020). Similarly to TSA-seq, TSA-MS exploits the predicted and measured exponential decay of tyramide (phenol)-biotin free radicals as they diffuse outwards from the immunostained target.

Given the ∼1-micron radius of this spreading (Chen et al., 2018), we realized that tyramide labeling of proteins would extend beyond the NS to proteins in a significant volume surrounding each NS. We therefore ranked proteins by their relative enrichment in NS versus near centromeres, stained by CENPA, to distinguish between proteins specifically enriched near NS versus proteins present throughout the nucleus, including near centromeres; we expected that centromeres would represent a nuclear local environment with low levels of proteins involved in gene expression.

Whereas 74% of the top 91 ranked proteins had been previously reported as colocalizing with NS, the next 92-200 top ranked proteins showed a sharp decrease in previously reported NS colocalization to ∼13% (Dopie et al., 2020). Despite this drop in NS colocalization, a large fraction of these proteins, similar to the fraction of known NS-enriched proteins, was connected to various steps in gene expression. Therefore, we reasoned these moderately-ranked proteins were likely enriched in transcriptionally active regions surrounding NS, and thus were candidates for proteins marking our hypothesized gene expression compartment(s) surrounding and extending outwards from NS (yellow regions in Fig. 1A).

We identified 25 candidate proteins from this list of moderately-enriched proteins that were reported to show non-diffuse immunostaining within the interchromatin space (ICS), based on the literature and/or the Human Protein Atlas (Pontén et al., 2009). As a first screen, we examined the nuclear distribution of each of these candidate proteins relative to both NS and DNA-staining, looking for candidate proteins that formed distinct staining patterns within and/or surrounding NS and extending out into areas of low DNA-staining corresponding to the ICS (Fig. 1B, left). We visualized their nuclear distribution either using transiently expressed EGFP fusion proteins or through immunostaining when antibodies were available.

Among these 25 proteins, essentially all showed some type of nondiffuse pattern, with foci of higher intensity throughout the nucleoplasm (SFig. 1A-B). However, there was a continuum in the extent of the nuclear area occupied outside of NS by higher than background intensities. Proteins such as SRRM2 and RBM25 showed little staining outside of NS, while proteins such as DDX41 and PNN showed staining occupying most of the nucleus (SFig. 1A-C). We focused on proteins which showed intermediate occupancy between these extremes, with patterns of staining extending between NS and throughout the nucleus, even extending all the way to the nuclear periphery (Fig. 1B, SFig. 1A-C).

For these proteins in this intermediate nuclear occupancy category (SFig. 1B-C), we observed two “perispeckle” categories of staining. We chose DDX39B and THRAP3 as founding members of these two intermediate patterns (SFig. 1C), which both occupied ∼40% of the nuclear area outside of NS (SFig. 1B). First, we observed proteins distributions similar to THRAP3 in which the protein was not concentrated within NS but was high in concentration surrounding NS (Fig. 1C, SFig. 1A). Second, we observed protein distributions similar to DDX39B in which the protein was most highly concentrated within NS (Fig. 1D, SFig.1A). These differences were made clearer by comparison of the cumulative protein intensities within versus outside NS (SFig. 1C). Finally, there were proteins that showed non-diffuse staining throughout the nuclear interior, but with little relationship to NS. Examples included SAFB and HNRNPM (SFig. 1A, 1C).

We decided to ask whether proteins within the same perispeckle categories might colocalize with each other. We did this by comparing protein colocalization with THRAP3 for the first category and DDX39B for the second category (Fig. 1B, right panel). To facilitate this screening, we used CRISPR-mediated knock-ins to fluorescently tag endogenously expressed proteins, constructing U2OS cell lines stably expressing either the combination of EGFP-SON and mCherry-THRAP3 or EGFP-SON and mCherry-DDX39B (see Methods). This avoided the overexpression induced changes in protein distribution sometimes seen after transient transfection (Fig. 1B).

We then used immunofluorescence to ask which other candidate proteins overlapped extensively with either DDX39B or THRAP3 (Fig. 1B). Due to the lack of a suitable antibody for MAGOH, we utilized lentivirus-based transduction of a mCherry-MAGOH expression cassette followed by immunostaining of THRAP3/DDX39B.

Several additional candidate proteins shared high colocalization for either THRAP3 or with DDX39B, suggesting that they were additional components of these perispeckle patterns. We named the intranuclear protein distribution overlapping with THRAP3 as “Perispeckle THRAP3 Pattern” (PSTP) and the intranuclear protein distribution overlapping with DDX39B as “Perispeckle DDX39B Pattern” (PSDP) (Fig. 1C-E). Additional proteins showed their own distinctive nuclear distributions (SFig. 1D); thus, the ICS is likely further subdivided into additional novel networks/condensates/bodies that might partially overlap with PSTP and/or PSDP. For example, PLRG1 was enriched inside NS, similar to PSDP, but outside of NS colocalized with PSTP (SFig. 1D, F).

PSTP markers currently include THRAP3 (BCLAF2) and its paralog BCLAF1 (Fig. 1 B-C,1E). Additionally, PLRG1 overlaps with PSTP outside of nuclear speckles (SFig. 1E). PSDP markers currently include DDX39B (UAP56), RBM8A (Y14), MAGOH, and POLDIP3 (Fig. 1D,F, SFig. 1D). DDX39B was previously described as enriched in NS but also enriched in the region surrounding NS (Kota et al., 2008). Here we show that DDX39B together with other PSDP components also radiate outwards from NS in thread-like projections that frequently connect neighboring speckles (Fig 1. D,F) while forming a network-like pattern which in some nuclear regions show small foci lining up in linear arrangements (Fig 1D-inset orange box).

Next, we examined how MALAT1 localized relative to these perispeckle patterns, as MALAT1-a long, noncoding RNA (lncRNA)- is spatially related to NS (Fei et al., 2017). Functionally, MALAT-1 associates with certain transcription factors and coactivators (Arun et al., 2020) and has been implicated in the regulation of alternative splicing (Miao et al., 2022). Using an oligolibrary containing 100 probes targeted against MALAT1, we observed both the high NS-enrichment of MALAT1 as well as single RNAs within the ICS. MALAT1 single RNAs away from NS were seen either overlapping or at the edge of both THRAP3 and DDX39B foci (SFig. 1G). Specifically, ∼50-60% of MALAT1 RNA signal within the interchromatin space overlapped with both THRAP3 and DDX39B outside NS, using Otsu thresholding (Otsu, 1979) to segment MALAT1, THRAP3, and DDX39B intensities (SFig. 1G).

In summary, we have identified two novel “perispeckle patterns” that subdivide the interchromatin space, as defined by the specific localization patterns of nuclear proteins already reported to be involved in gene expression. Moving forward, we focused the differential localization of THRAP3 (PSTP) versus DDX39B (PSDP) as markers of these two perispeckle staining distributions.

### Highly dynamic PSTP and PSDP networks which repeatedly recur within the same or nearby interchromatin space

Projections of higher intensity THRAP3 and DDX39B staining extend outwards from NS forming apparent networks within the interchromatin space.

Fixation can alter the cellular distribution of intrinsically disordered domain-containing proteins (Irgen-Gioro et al., 2022). We therefore used live-cell imaging in U2OS cells to confirm similar PSTP and PSDP distributions in live cells and visualize their dynamics over seconds (Fig. 2A-B) to hours (Fig. 2C-D) using the dual SON and THRAP3 or SON and DDX39B tagged cell lines.

**Figure 2:**
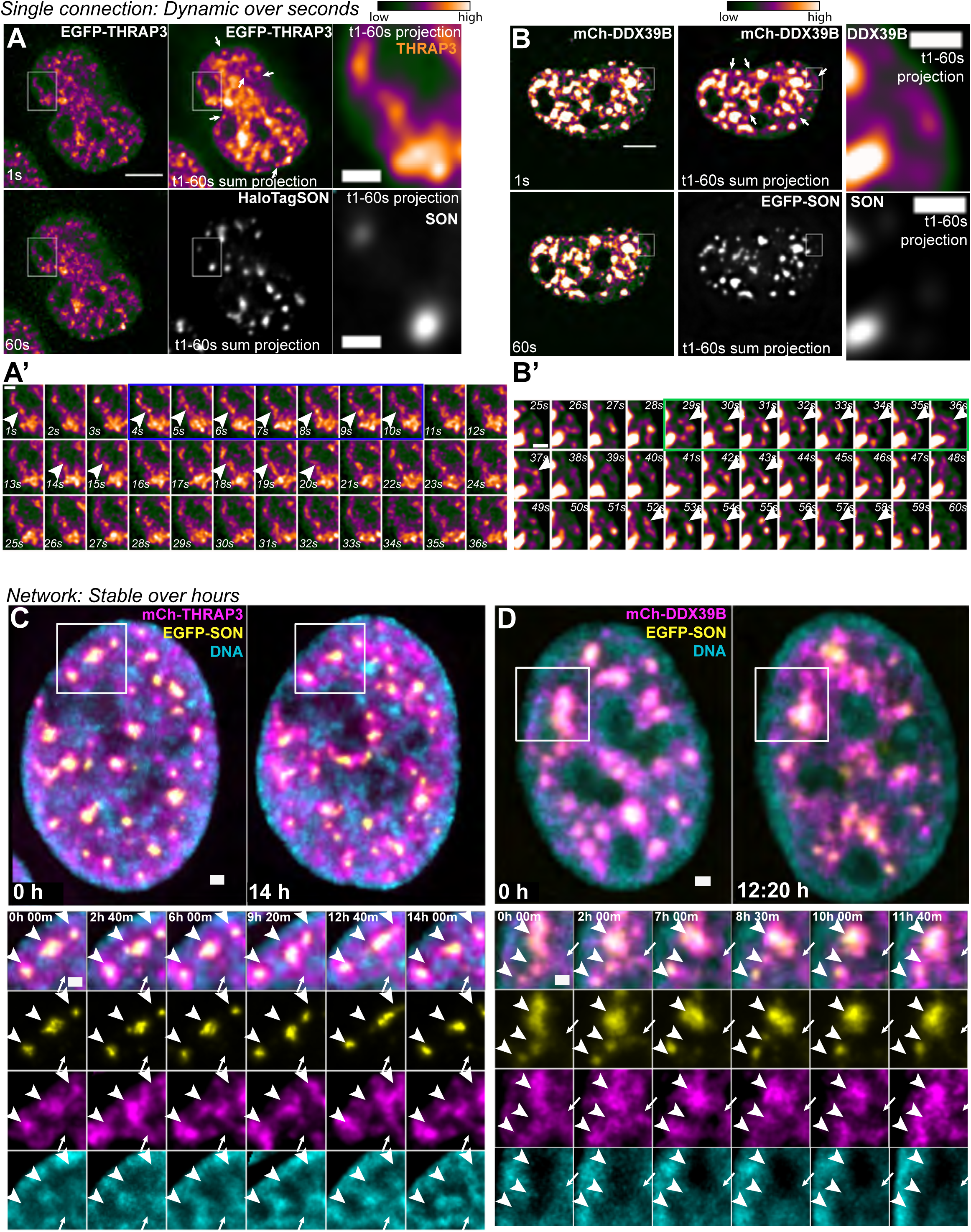
Highly dynamic PSTP and PSDP networks appear, disappear, and reappear within the same or nearby IC space. A and B: Live-cell imaging of endogenously tagged EGFP-THRAP3 (A, Video 1) and mCherry-DDX39B (B, Video 2) along with SON (gray) over 1 second (s) intervals shows linear thread-like protein concentrations (“structures”) appear and disappear within the vicinity of NS (see Videos 1 and 2). Time projections over 60 seconds (middle and right panels) show these thread-like structures more clearly, indicating they consistently recur within the same spatial locations, with some foci (arrows) showing a relative increase in brightness, suggesting more stable and/or frequent accumulation of protein at these regions over time. Scale bar=5 μm. Insets (A’ and B’) show the dynamics of single thread-like structures over 1 second intervals. Blue (A’) and green (B’) boxes show sequential frames in which thread-like structures form and remain stable over a number of seconds. These structures then disappear but reform at later times (arrowheads) in the same region. Sixty second time projection of the same structures are shown enlarged in insets ((A)and (B), right). Scale bar= 1 μm. THRAP3 and DDX39B intensities are scaled using a blue-orange LUT (Fiji, blue-orange-icb LUT). C and D: Endogenously tagged mCherry-THRAP3 (C; magenta, Video 3) and mCherry-DDX39B (D; magenta, Video 4) networks are stable within the interchromatin (IC) space over hours (∼10-14 hours (h)) in live cells (see Videos 3 and 4). Insets (bottom panels) show boxed region (top panel, white borders) at multiple time-points. Nuclear speckles (yellow) form and fuse and/or remain stable (arrowheads) within the vicinity of perispeckle networks, while thread-like projections of THRAP3 or DDX39B appear, disappear, and then reappear repeatedly within the same regions (arrows). DNA (cyan) is visualized by SiR-Hoechst staining. Scale bar= 1 μm.

Both THRAP3 and DDX39B show non-diffuse high local concentration in linear extensions that appear and disappear every few seconds in the vicinity of speckles (Fig. 2A&A’, 2B&B’, Videos 1&2). Summing these 2D projections over a time of 1 minute reveals a clearer network distribution for both THRAP3 and DDX39B due to repeated recurrence of these projections in the same spatial locations (Fig. 2A-B, white arrows). Thus, while both the PSTP and PSDP are highly dynamic in a timescale of seconds, their extensions averaged over time are quasi-stable over a timescale of minutes.

Over a much longer timescale of up to 10 hours, PSTP and PSDP networks preserve their overall nuclear distributions despite their fast dynamics (Fig. 2C-D). NS show occasional movements and fusion and fission events within relatively stable DNA-depleted regions (Videos 3&4). PSTP and PSDP were more dynamic, showing repeated appearance/disappearance of thread-like extensions over several minute intervals within the same DNA-depleted spaces surrounding and between NS (Fig. 2C&D Videos 3&4). Because these projections tend to reform in the same or similar regions over time, the overall nuclear distributions of PSTP and PSDP remain relatively stable over longer time periods. In a companion paper, we explore these dynamics in greater detail for both SON and two PSDP members-DDX39B and MAGOH (Kim et al., 2025).

### PSTP and PSDP subdivide the interchromatin space and respond differently to functional and physiological perturbations

THRAP3 and DDX39B, our marker proteins for PSTP and PSDP respectively, were both previously reported as overlapping with and/or surrounding NS (Lee et al., 2010; Kota et al., 2008). Because PSTP and PSDP markers also occupy interchromatin space further away from NS and throughout the nucleus, we next examined their relative spatial distribution outside of NS. Across the range of spatial resolution provided by wide-field deconvolution and STORM super resolution light microscopy, THRAP3 and DDX39B staining “networks” appear largely juxtaposed rather than coincident within the interchromatin space outside of NS (Fig. 3A-B). They are significantly less colocalized relative to each other than the colocalization seen for other components of PSTP with THRAP3 or other components of PSDP with DDX39B (Fig.1E-F, SFig.1F-G).

**Figure 3:**
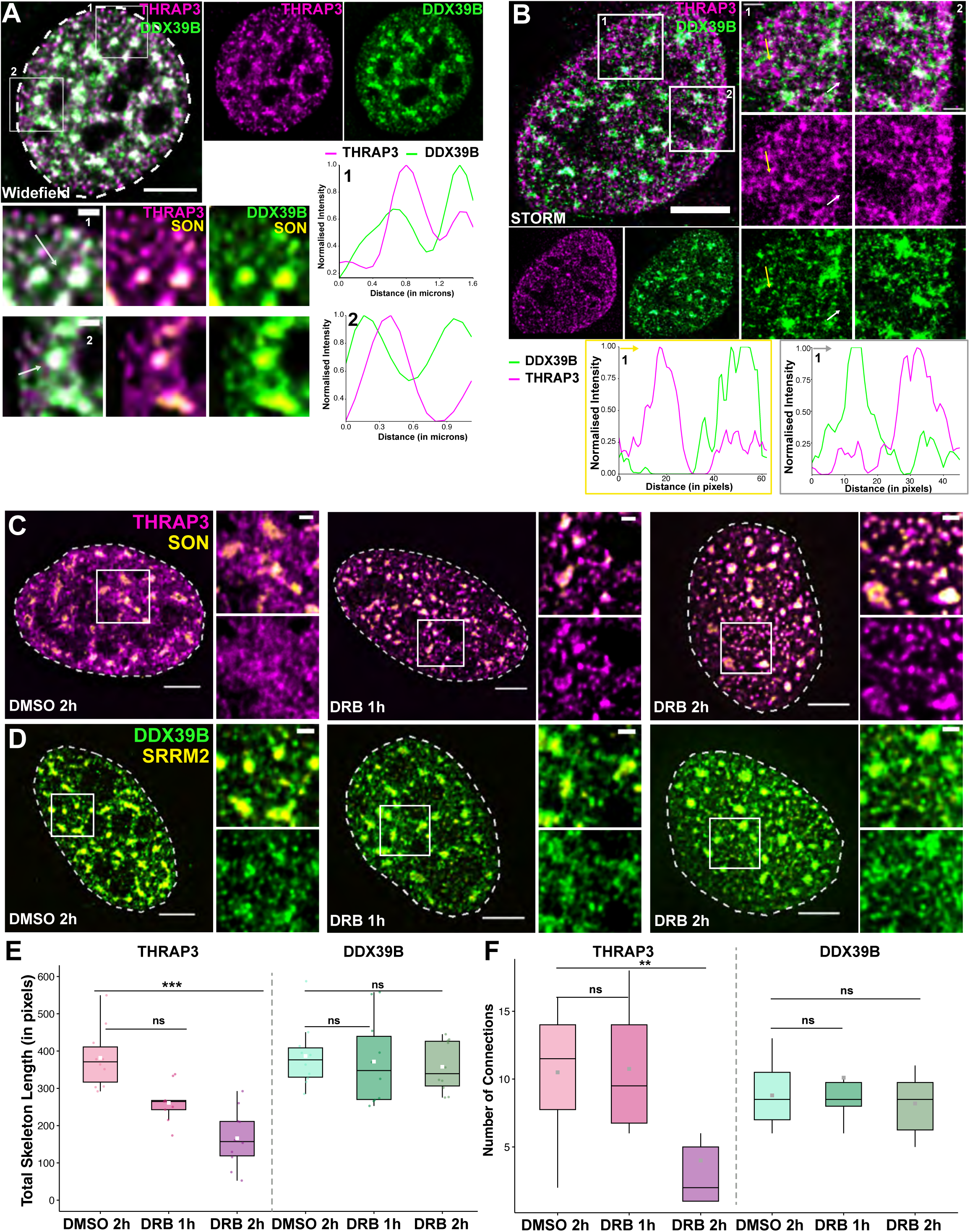
PSTP and PSDP subdivide the interchromatin space and respond differently to functional perturbation. A: Wide-field imaging of immunostained THRAP3 (magenta) and DDX39B (green) relative to SON (EGFP-SON knock-in; yellow) in U2OS cells shows that the two perispeckle networks are juxtaposed outside of NS. Insets show enlargements of boxed regions (rectangles 1,2). Line profile plots (along white arrows within insets) show THRAP3 and DDX39B peaks are offset from each other. Scale bars-5 μm main image; 1 μm inset. B: STORM super-resolution imaging of immunostained THRAP3 (magenta) and DDX39B (green) demonstrates more clearly the spatial separation of THRAP3 versus DDX39B concentrations suggested by wide-field microscopy (A). Insets are enlargements of boxed regions (white rectangles 1, 2) and highlight nuclear regions showing THRAP3 and DDX39B occupying distinct spaces. Line profile plots (along yellow or white arrows within inset 1) show peaks of THRAP3 and DDX39B concentrations separated by distances of ∼20-30 nm. Scale bars-5 μm main image; 1 μm inset. C-D: Transcriptional inhibition by 5,6-Dichloro-1-β-D-ribofuranosylbenzimidazole (DRB) treatment disrupts THRAP3 perispeckle pattern (C; magenta) but DDX39B perispeckle pattern (D; green) remains relatively unchanged. NS (yellow) are marked by SON (C-top) or SRRM2 (D-bottom). THRAP3 progressively condenses into distinct foci arranged frequently in thread-like structures and ultimately accumulates in cap-like accumulations adjacent to NS (Video 5). Scale bars-5 μm main image; 1 μm inset. E-F: Box plots showing total skeleton length (E) and number of connections (F) of THRAP3 and DDX39B networks (SFig. 3A) upon DRB treatment (1, 2 hours (h)) as compared to DMSO (2 hours). Central line indicates median and square within the boxplot indicates mean. n (number of nuclei) = 10 for each condition. One-way ANOVA with Tukey post-hoc test was used to calculate P-values. ns : P > 0.05; ** : P ≤ 0.01; ***: P ≤ 0.001.

After transcriptional inhibition, many NS components, including SON and a number of splicing factors, are known to concentrate further within NS with a concomitant decrease in their nucleoplasmic concentrations (Spector and Lamond, 2011); this is accompanied by NS rounding, enlargement and fusion (Kim et al., 2019).

However, the THRAP3 and DDX39B staining networks responded differently after transcriptional inhibition, providing further support for their independent nuclear localization. Between 0-2 hours of transcriptional inhibition by DRB (5,6-dichloro-1-beta-D-ribofuranosylbenzimidazole), the thread-like, diffuse network of THRAP3 staining outside of NS consolidates into fewer but more distinct threads, that then breakup into distinct foci aligned along linear paths, plus larger concentrations adjacent to and surrounding NS (Fig. 3C, Video 5). In contrast, the DDX39B distribution appeared relatively intact, although their stability over time may decrease as reported elsewhere (Kim et al., 2025) (Fig. 3D). DRB inhibits CDK9 and thus transcriptional pause release (Bensaude, 2011; Yankulov et al., 1995). We observed similar disruption but to a lesser extent after inhibiting transcription instead using either Triptolide (SFig 3A, which inhibits the XBP subunit of TFIIH and thus transcriptional initiation (Titov et al., 2011), and α-amanitin (SFig. 3B), which initially inhibits but then degrades RNA Pol II over the ∼6 hr treatment time (Nguyen, 1996).

To quantify the changes in THRAP3 network upon DRB treatment, we used a function “skeletonize” which uses binary thinning/erosion until a one-pixel wide representation remains. These one-pixel wide chains are called “skeletons” that retain both the shape and connectivity of the original image (SFig. 3C). Using this skeletonize function, we measured two parameters per nucleus: 1) total skeleton length, defined as the sum of all skeletons; 2) the number of connections, where each connection is defined as group of connected pixels >5 pixel length.

Based on these parameters, the THRAP3 distribution shows both a reduction in total skeleton length and in the number of connections. In contrast, the DDX39B distribution remains relatively intact (Fig. 3E-F). DRB treatment similarly disrupted PSTP member BCLAF1 distribution which also formed foci which colocalized with THRAP3 foci (SFig. 3D-E). However, PSDP member RBM8A shows a heterogenous response within the cell population, ranging from minimal to relatively significant perturbation (SFig.3F-G). The RBM8A network, on average, breaks up from a continuous structure to multiple long thread-like structures with DRB treatment as reflected in both the decrease in total skeleton length and increase in the number of connections (SFig. 3G-H).

In summary, we conclude that PSTP and PSDP form spatially and functionally distinct networks that subdivide the interchromatin space.

### RNA Pol II Ser2p foci localize in close proximity to the PSTP and PSDP networks

We next examined the spatial distribution of RNA Pol II foci relative to PSTP and PSDP (Fig. 4). Antibodies against the Ser5p and Ser2p epitopes of the C-terminal domain (CTD) of the largest RNA Pol II subunit recognize the early initiation versus later elongating forms of RNA Pol II, respectively (reviewed in (Egloff and Murphy, 2008)). Immunostaining using these antibodies reveals ∼60 nm diameter foci, originally termed “RNA Pol II factories” that map near sites of transcription (Jackson et al., 1993; Iborra et al., 1996; Pombo, 1999). Initially proposed as actual transcription sites (Martin and Pombo, 2003; Xie et al., 2006), the functional significance of these transcription factories remains unclear.

**Figure 4:**
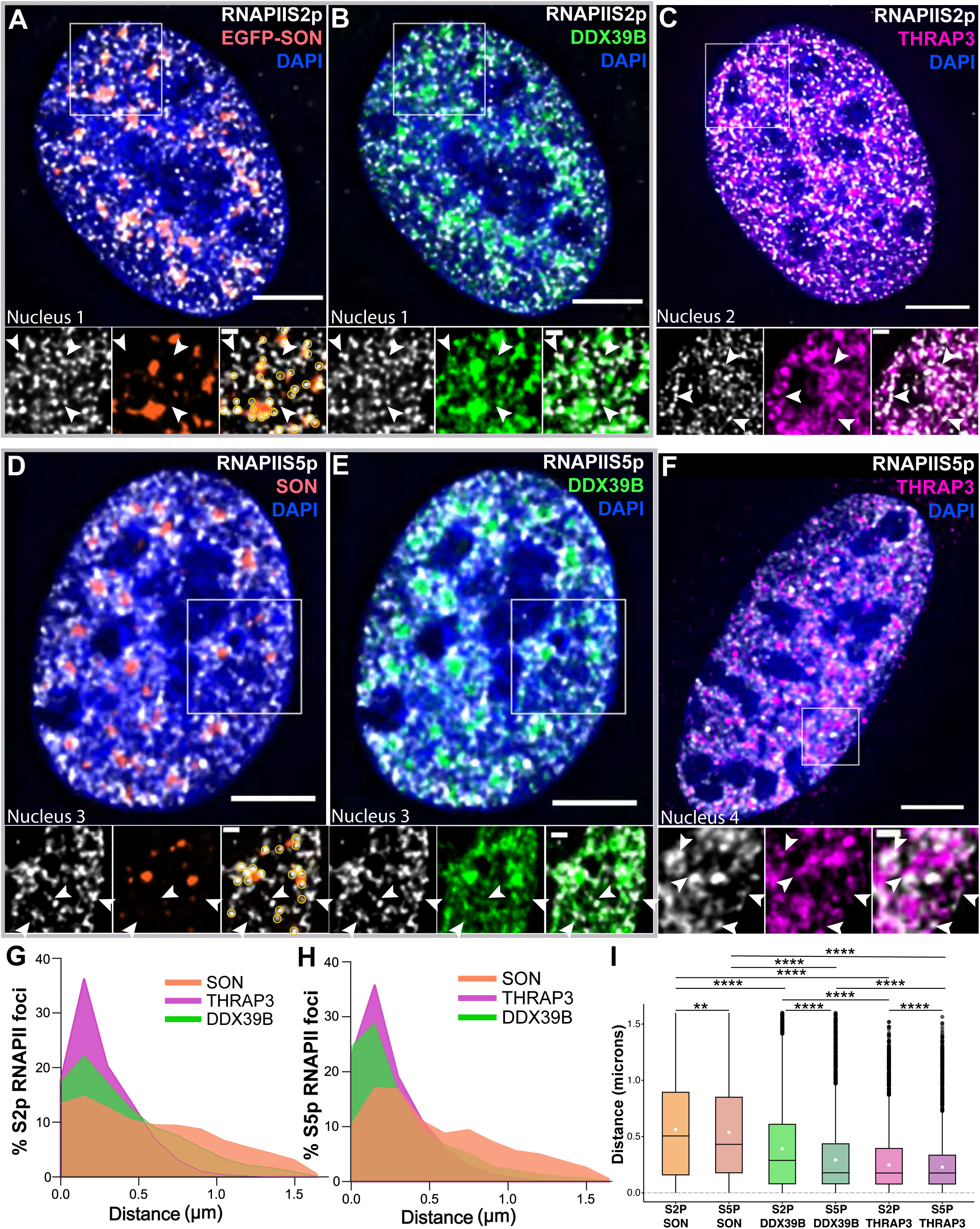
RNA pol2 Ser2p foci localize in close proximity to PSTP and PSDP. A-F: Immunostaining of RNA Pol II serine 2 phosphorylated foci (white) (A-C) and serine 5 phosphorylated foci (white) (D-F): RNA pol II foci (arrowheads) not near NS (SON, orange; endogenously tagged with EGFP) align adjacent to THRAP3 (magenta; immunostaining) and DDX39B (green; endogenously tagged with mCherry) foci within the IC space. RNA Pol II foci at the edge of SON-marked speckles indicated by yellow circles. Images A-B and D-E (gray boxes) show Pol II foci in the same nucleus relative to SON and DDX39B. G-H Distance distribution curves for the percentage of RNA Pol II serine 2 phosphorylated foci (G) and serine 5 phosphorylated foci (H) relative to SON, THRAP3 and DDX39B. I: Boxplots showing average distance of foci relative to RNA Pol II foci relative to SON, THRAP3 and DDX39B. Central line indicates median distance and square within the boxplot indicates mean distance. P-values calculated using Wilcoxson sum rank test; **: P ≤ 0.01; ****: P ≤ 0.0001. (n=9124-81267 RNAPII foci).

Although an increased density of RNA Pol II Ser5p and Ser2p foci have previously been described as surrounding the NS periphery, large numbers of RNA Pol II Ser2p and Ser5p foci away from NS are distributed throughout the nuclear interior in no obvious pattern (Chen et al., 2018) (Fig. 4A&D). Immunostaining, however, now reveals a striking spatial correlation of these RNA Pol II foci localized away from NS with the THRAP3 and DDX39B networks (Fig. 4B-C and 4E-F).

We used an intensity threshold of 40% of local intensity maxima to segment foci of both THRAP3 and DDX39B (SFig. 4). Median distances of both Ser5p and Ser2p RNA Pol II foci are ∼0.2 μm to the nearest edges of THRAP3 foci (Fig. 4K, magenta and purple). Ser5p RNA Pol II foci are similarly close to DDX39B foci (Fig. 4K, light green), while Ser2p RNA Pol II foci show a somewhat increased distance distribution but remain mostly within ∼0.5 μm to DDX39B foci (Fig. 4K, dark green).

Thus, RNA Pol II foci, used as proxies for mapping sites of active transcription, nearly all map within a few hundred nm of THRAP3 or DDX39B foci, supporting the idea of PSTP and PSDP acting as additional nuclear niches closely associated with transcriptionally active chromosome loci.

### Gene expression hot zones interact preferentially with PSTP and PSDP as measured by TSA-seq

To follow up our demonstration of the close association of RNA pol II foci with perispeckle patterns, we next asked if, as hypothesized (Fig. 1A), Type II SON TSA-seq peaks would show a close association with one or both perispeckle patterns similar to how Type I SON TSA-seq peaks show a close association with NS. More specifically, we used TSA-seq to compare the association of Type II SON TSA-seq peaks with the PSTP and/or the PSDP versus NS.

We combined the “super-saturation” TSA-seq 2.0 protocol (Zhang et al., 2021) with the use of dithiothreitol (DTT) to improve the TSA-seq spatial resolution, as described in the original TSA-seq 1.0 protocol (Chen et al., 2018). DTT improves spatial resolution by decreasing the lifetime, and therefore the diffusion radius of the tyramide free-radical, while the “super-saturation” protocol greatly reduces the number of cells required for TSA-seq 2.0. To compensate for the reduced labelling due to quenching by DTT, we increased the tyramide-biotin concentration by 10-fold.

Both in HCT116 (Fig. 5A-B) and U2OS cells (SFig. 5A-B), DDX39B and THRAP3 TSA-seq peak and valley patterns qualitatively roughly parallel the SON TSA-seq peak and valley pattern. Quantitatively, however, there are informative differences. We interpret DDX39B and THRAP3 TSA-seq peaks as corresponding to genomic regions that associate closest to foci of DDX39B and THRAP3 staining. The highest amplitude DDX39B and THRAP3 peaks do align with the largest, Type I SON TSA-seq peaks (yellow highlights, Fig. 5A, SFig. 5A). However, we interpret the disproportionally higher amplitude of DDX39B and THRAP3 relative to smaller SON TSA-seq peaks as evidence for the association of these genomic regions near foci of DDX39B and THRAP3 when they are not located near NS. For example, DDX39B and THRAP3 TSA-seq peak amplitudes are nearly the same over Type I SON TSA-seq peaks that vary two-fold in SON TSA-seq amplitudes (yellow highlights, Fig. 5A, SFig. 5A). As hypothesized, Type II SON TSA-seq peaks with near-zero SON TSA-seq amplitudes show disproportionally larger DDX39B and, especially, THRAP3 TSA-seq peak amplitudes (blue highlights, Fig. 5A, SFig. 5A) that in some cases approach similar THRAP3 peak amplitudes as observed over Type I peaks (yellow highlights, Fig. 5A, SFig 5A). This disproportionate scaling extends more generally over larger regions of the genome as appreciated in scatterplots of DDX39B or THRAP3 versus SON TSA-seq which show a lower slope for higher versus lower values of SON TSA-seq (Fig. 5B).

**Figure 5:**
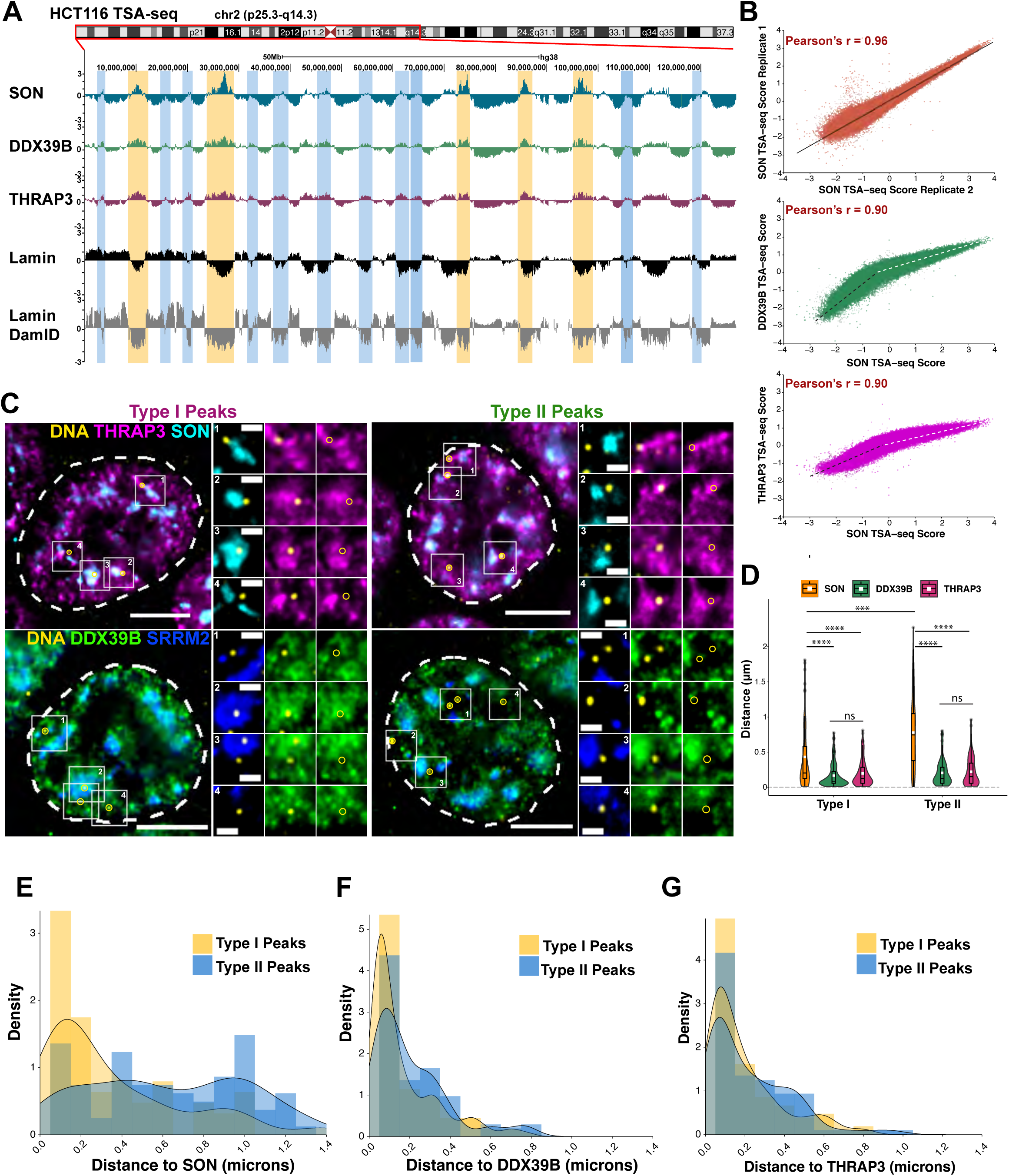
Gene expression hot-zones interact preferentially with PSTP and PSDP in HCT116 cells. A: Representative TSA-seq plot for chromosome 2 in HCT116 cells for SON (blue), DDX39B (green), THRAP3 (magenta), Lamin B1 (black), and Lamin B1 Dam-ID (gray). Yellow and blue highlights mark Type I peaks and Type II peaks, respectively. Type I peaks show large magnitude peaks for SON but lower amplitude and broader peaks for DDX39B and THRAP3. Type II peaks show well-defined and higher magnitude peaks for DDX39B and THRAP3 as compared to SON. B: Scatterplots between two SON TSA-seq replicates (orange), SON vs DDX39B (green) and SON vs THRAP3 (magenta). Overall, TSA-seq scores for DDX39B and THRAP3 correlate closely with SON. Both DDX39B and THRAP3 versus SON scatterplots show two regions with distinct slopes: higher for negative SON scores (black dotted line) but lower for positive SON scores (white dotted line). C: DNA-FISH (yellow) in HCT116 cells for multiple Type I and Type II peaks (apex) shows higher frequency of association with NS (SON-cyan or SRRM2-blue) for Type I versus Type II peaks. NS non-associated alleles show close proximity or locate adjacent to THRAP3 (magenta) or DDX39B (green) foci/accumulations. Regions highlighted in white boxes are shown on the right as insets. Scale bars-5 μm main image; 1 μm inset. D: Violin plots showing distance of DNA-FISH spots relative to SON, DDX39B and THRAP3 foci/accumulations. Central line indicates median distance and white square within the boxplot indicates mean distance. (n (alleles) as indicated: 67 (Type I) and 95 (Type II) SON; 114 (Type I) and 103 (Type II) DDX39B; 105 (Type I) and 96 (Type II) THRAP3). One-way ANOVA with Tukey post-hoc test was used to calculate P-values. ns : P > 0.05; *** : P ≤ 0.001; **** : P ≤ 0.0001. E-G: Probability density plots for distance distribution of DNA-FISH spots relative to SON (E), DDX39B (F) and THRAP3 (G) foci/accumulations for Type I (yellow) and Type II (blue) peaks. (n (alleles) as indicated: 67 (Type I) and 95 (Type II) SON; 114 (Type I) and 103 (Type II) DDX39B; 105 (Type I) and 96 (Type II) THRAP3).

Finally, DDX39B and THRAP3 peaks are significantly broader and more rounded than their corresponding SON TSA-seq peaks, and they extend over most of the iLADs in which they are embedded, in contrast to the narrower SON TSA-seq peaks. This trend is somewhat more pronounced in the THRAP3 versus DDX39B TSA-seq.

We interpret this broader shape of DDX39B and THRAP3 TSA-seq peaks relative to SON TSA-seq peaks as due to the extent to which DDX39B and THRAP3 networks extend throughout the nuclear interior such that iLAD regions are never that distant from the nearest DDX39B or THRAP3 foci. The even broader shape of the THRAP3 versus DDX39B TSA-seq peaks would then be consistent with the greater extent to which the THRAP3 network spreads throughout the nuclear interior relative to the DDX39B network.

### FISH validation of close association of gene expression hot-zones with PSTP and PSDP

To test these TSA-seq interpretations, we first selected 7 large SON TSA-seq Type I peaks and 8 Type II SON TSA-seq local maxima (small peaks or “peak-within-valleys”) for immuno-FISH validation in both HCT116 (Fig. 5C) and U2OS (SFig. 5C) cells. We used pooled oligonucleotide probes that targeted simultaneously either all 7 Type I peaks or all 8 Type II SON TSA-seq peaks (Supplementary Table 1). To determine distances to the nearest NS, defined by SON staining, or to the nearest DDX39B or THRAP3 foci, we used a local segmentation of the immunostaining based on an intensity threshold corresponding to 50% of the peak intensity of the nearest staining focus (see Methods).

As expected, the aggregated Type I SON TSA-seq peaks localized near NS (Fig. 5C-D, SFig. 5C-D), with median distances of 0.20 (HCT116) and <0.20 (U2OS) μm from NS. The aggregated Type II peaks instead localized at varying distances from NS, with median distances of 0.74 (HCT116) and 0.54 (U2OS) μm. In contrast, in both HCT116 and U2OS cells Type I and II SON TSA-seq peaks positioned close to THRAP3 or DDX39B foci, with median distances of ≤ 0.25 μm, including alleles that are positioned away from NS (Fig. 5B-G, SFig. 5C-G).

We next used DNA FISH in U2OS cells to paint chromosome “trajectories” from the nearest LAD/iLAD boundary to either the apex of a SON TSA-seq Type I peak or to the apex of a SON TSA-seq Type II peak (Fig. 6). In both cases, we used oligonucleotide libraries designed to target 7-9 ∼100 kb regions spaced approximately evenly along these trajectories.

**Figure 6:**
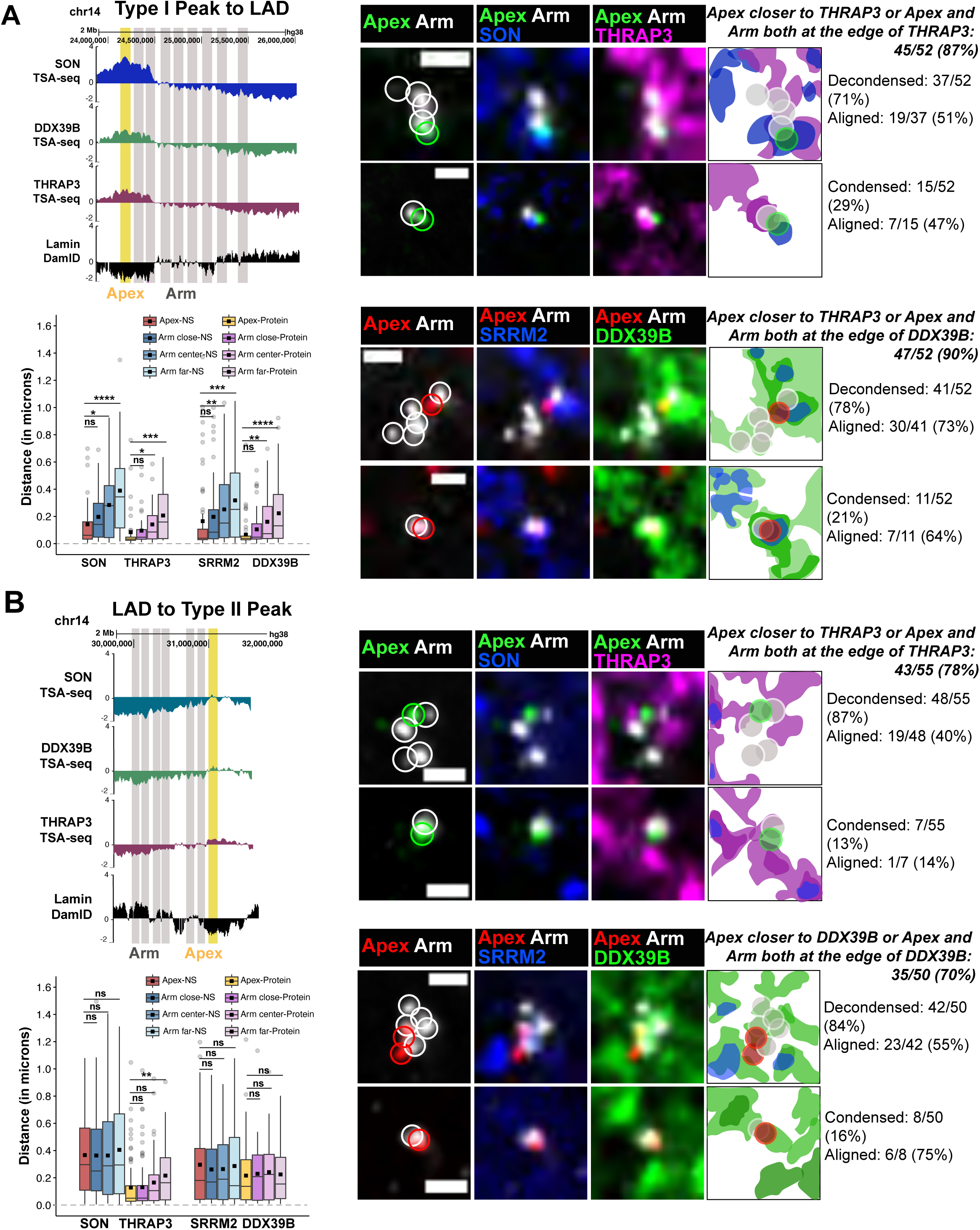
Mapping of chromosome trajectory-SON TSA-seq peak to nearest LAD-shows alignment along PSTP and PSDP foci / concentrations. (A-B) Left, top panels: Browser views showing SON, DDX39B, THRAP3, Lamin B1 TSA-seq and Lamin B1 DamID tracks corresponding to one Type I (A) or Type 2 (B) SON TSA-seq peak-to-valley chromosome trajectories mapped by DNA FISH. Highlights show locations of peak “Apex” (yellow) and multiple “Arm” (grey) DNA FISH probes. (A-B) Right: Sum-intensity projections of Type I (A) or Type II (B) Apex versus Arm probes along peak-to-valley chromosome trajectories and their spatial positioning relative to THRAP3 (magenta), DDX39B (green), and nuclear speckles (SON/SRRM2; blue) immunostaining. The percentage of DNA trajectories appearing aligned with perispeckle staining is higher for DDX39B versus THRAP3. Scalebar=1 μm. (A-B) Left, bottom panels: Statistics for Apex versus Arm chromosome orientation relative to foci of SON, THRAP3, and DDX39B summarized in box plots. Apex FISH signals are statistically significantly closer to SON, THRAP3, and DDX39B foci than to middle and distal Arm FISH signals for the Type I trajectory. Apex FISH signals are closer to middle and statistically significantly closer to distal Arm signals relative to THRAP3 foci, but not SON or DDX39B foci, consistent with the distal positioning of Type II trajectories relative to NS and the higher trajectory alignment seen relative to the DDX39B versus THRAP3 staining. Black squares represent mean distances and central line represents median distances. n (number of trajectories analyzed): Type I trajectory: THRAP3/SON = 40; DDX39B/SRRM2= 66; Type II trajectory: THRAP3/SON=82; DDX39B/SRRM2=54. Kruskal-Wallis test followed by Dunn’s post-hoc test was used to calculate P-values. ns : P > 0.05; * : P ≤ 0.05; ** : P ≤ 0.01; *** : P ≤ 0.001; **** : P ≤ 0.0001.

In most cells (∼70-90%), both chromosome trajectories showed multiple, individual DNA FISH spots corresponding to the iLAD regions flanking the TSA-seq peak locations, consistent with a partly to more fully extended iLAD chromosome trajectory (Fig. 6). In some cells (∼10-30%), however, these FISH spots merged into a single, compact signal, larger than the wide-field diffraction limit, suggesting a more localized folding of the iLAD trajectory (Fig. 6). Interestingly, a high fraction of the extended trajectories aligned relative to the THRAP3 (40-51%) and, especially, the DDX39B staining foci (55-73%) (Fig. 6).

The arms of both extended and compact trajectories appeared nearly always (∼70-90%) as close or further from the nearest SON, THRAP3, or DDX39B foci than the DNA FISH spot corresponding to the apex of the TSA-seq peak location (“Apex”) (Fig. 6).

To compare quantitatively the association of different trajectory regions with the perispeckle patterns and NS, we segmented and divided the DNA-FISH arm signal into three parts based on their proximity to the Apex: 1) “Arm-close” region (pixels with distances ≤25^th^ percentile closest to Apex); 2) “Arm-center” region (pixels with distances within the 26th-75^th^ percentile from Apex); and 3) “Arm-far” region (pixels with distances >75^th^ percentile from Apex). Then for the Apex and each of these three arm regions, we calculated the shortest Euclidian distance of the center of mass to the nearest segmented NS or protein foci edge. (Note that the nearest NS or protein foci edge could be different for the Apex and each of these three regions; see Methods.)

For the Type I trajectory, the Apex was closer to SON, THRAP3 or DDX39B foci compared to Arm regions (Fig. 6A, left lower panel). Moreover, the “Arm-close”, “Arm-center”, and “Arm-far” regions showed a progressive increase in distances relative to the nearest NS or protein foci edge, respectively. As predicted from the rounded shape of the DDX39B and THRAP3 TSA-seq Type II peaks, the entire Type II iLAD chromosome trajectory typically lies within several hundred nm of some part of the DDX39B and THRAP3 network (Fig. 6B). Distances of the Apex of the Type II peak from the nearest NS edge as expected were larger than for the Type I peak (∼0.4 versus <0.2 μm). The Type II Apex compared to the Arm-far region was significantly closer to the THRAP3 foci (Fig. 6B, lower left panel). However, the distances to the edge of the nearest DDX39B foci were similar for the Apex and all three arm regions (Fig. 6B, lower left panel), which might reflect the apparent higher alignment of the Type II overall trajectory with the DDX39B network noted previously.

In summary, these results confirm that the chromosome gene expression hot-zones corresponding to SON TSA-seq local maxima associate close to DDX39B and THRAP3 foci, even when they are positioned at a distance from NS.

### Interactions of specific RNAs with PSTP and PSDP

NS were first proposed to act as gene expression hubs not only because a subset of active genes closely associated with the NS periphery but also because a subset of the RNAs transcribed from these genes, for example from the *COL1A1* and *actin* genes, specifically entered and concentrated within the adjacent NS (Hall et al., 2006). We later described how HSPA1A RNA also entered and concentrated within NS (Hu et al., 2009). As reviewed elsewhere, the trafficking of a large number of specific RNAs through NS has now been suggested using genome-wide sequencing-based methods (Wu et al., 2024; Barutcu et al., 2022; Khyzha et al., 2025). Most NS-enriched factors are involved in some aspect of RNA processing and/or export, and the idea of NS acting as a gene expression hub has always included the concept that proximity of genes and RNAs to NS might increase overall gene expression levels largely through their acceleration of post-transcriptional RNA processing and export events (Smith et al., 2020; Bhat et al., 2024; Barutcu et al., 2022; Wu et al., 2024; Wang et al., 2018).

The marker proteins for PSTP and PSDP that we identified here have similarly been functionally tied to various aspects of RNA processing, stability, and export. Therefore, using two experimental systems we decided to explore whether PSTP and/or PSDP networks might not only be associated with specific chromosome regions but also with specific pre-mRNAs and mRNAs, similarly to NS. Because Type I versus Type II chromosome regions are enriched in genes with low versus high intron-content, respectively (Gholamalamdari et al., 2025), we specifically compared the localization of RNAs from intronless versus intron-containing genes.

First, we used two cell lines, described elsewhere (Mor et al., 2010; Brody et al., 2011), carrying different stably integrated, multi-copy transgene arrays. Because these transgene arrays express at high levels, they produce large numbers of RNPs whose locations can be mapped relative to NS and perispeckle patterns with high statistical power within individual nuclei. The “E6” cell line contains a synthetic transgene with β-globin derived sequences including 6 exons and 5 introns, whereas the “Mini-dystrophin” cell line (“mini-Dys”) contains a truncated dystrophin transgene with a single exon and no introns. Both transgene RNAs contain 24 MS2 repeats and both cell lines express the MS2 coat protein (MCP) tagged with YFP that binds to this MS2 RNA repeat. Because of the hundreds of transcripts per nucleus, we used an automated image processing pipeline that segments the SON, DDX39B or THRAP3 intensities, using Otsu global thresholding, and then computes the distance of the center of each detected RNP to the nearest segmented staining feature (see Methods).

As our second experimental system, we used HSPA1 and HSPH1 endogenous genes, thereby allowing us to complement our results using the transgene arrays. Unlike most endogenous genes, HSPA1 and HSPH1 genes produce a high number of transcripts. We chose times after heat shock (HS) producing high numbers of HSPA1 and HSPH1 transcripts per nucleus, allowing us to avoid the use of transgenes while still acquiring measurements for ∼10-20 RNPs per nucleus away from the site of transcription. Whereas the highly homologous *HSPA1A* and *HSPA1B* genes expressed from the HSPA1 locus (situated within a SON Type I peak) produce short transcripts less than 1kb with no introns, the 27 kb *HSPH1* gene (resident within a SON Type II peak) contains 17 introns. The smaller number of transcripts per nucleus of these heat shock genes allowed us to use a semi-automated local intensity thresholding and measure distance of each RNP to the nearest staining foci by an automated algorithm (see Methods). This local thresholding allows detection of lower intensity staining foci which are missed by global thresholding approaches. The short length of HSPA1 genes results in significantly faster induction after HS as compared to the much longer HSPH1 gene. We chose different times after HS-30 mins for HSPA1 and 60 mins for HSPH1, which maximized the number of nuclear RNPs present in the nucleus away from their transcription site for each gene.

In both experimental systems, the intron-rich E6 and HSPH1 RNAs localized very close to THRAP3 foci in absolute distances (0.12 and 0.03 μm mean and median (E6); 0.13 and 0.07 μm mean and median (HSPH1)) and closer than the localization to THRAP3 foci of the intronless mini-dystrophin and HSPA1 RNAs (0.30 and 0.23 μm mean and median (mini-dystrophin); 0.34 and 0.30 μm mean and median (HSPA1)) (Fig. 7A-B,E&G, I-J). However, both intron-rich and intronless transcripts showed very close and similar distances to the nearest DDX39B foci (0.10 and 0.04 μm mean and median (E6); 0.17 and 0.06 μm mean and median (HSPH1); 0.11 and 0.05 μm mean and median (mini-dystrophin); 0.15 and 0.06 μm mean and median (HSPA1)) (Fig. 7C-D, F&H, I-J). These distances to the nearest DDX39B and THRAP3 foci for all these transcripts are significantly lower than their distances to NS (SON and SRRM2); this indicates that a significant fraction of the transcripts associate with DDX39B and THRAP3 foci away from NS.

**Figure 7:**
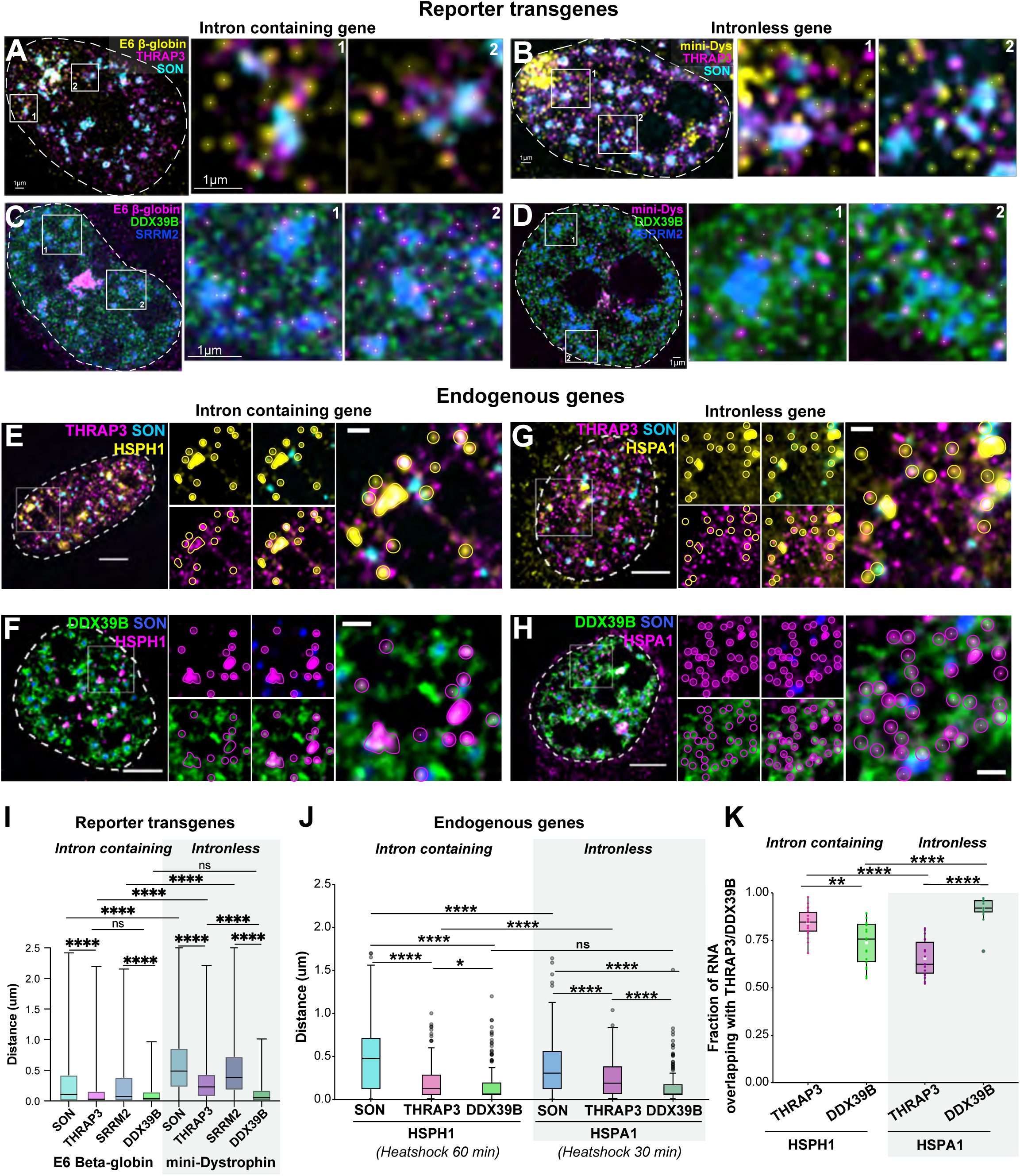
RNA from intron-containing and intronless genes associate differentially with PSTP and PSDP. A-B: RNAs (yellow) produced by the transgene array expressing the “E6” (6 exons) construct with βeta-globin sequences (A) localize more closely to THRAP3 foci / concentrations (magenta) than RNAs produced by the transgene array expressing the “mini-Dystrophin” (B) construct with a single exon (no introns). SON (NS) is cyan. Both RNAs are tagged with MS2 repeats and visualized by expression of the MS2-coat protein fused with YFP. (C-D) RNAs produced by both the E6 (C) and mini-Dystrophin (D) constructs localize closely with DDX39B (green) foci / concentrations. SRMM2 (NS) is blue. (E-F) FISH signals (yellow) for RNAs produced by the endogenous HSPH1 gene with many introns (E) localize closer to THRAP3 foci / concentrations (magenta) than RNAs produced by the endogenous HSPA1A and HSPA1B genes with no introns (F). SON (NS) is cyan. (G-H) FISH signals (magenta) for RNAs produced by the endogenous HSPH1 gene with many introns (E) and the endogenous HSPA1A and HSPA1B genes with no introns (H) both localize close to DDX39B (green) SRRM2 (NS) is blue. I: Box-plots of distances of β-globin E6 and mini-Dystrophin RNAs relative to SON, THRAP3, SRRM2 and DDX39B. n (number of RNPs per sample)= 1212-2627. Kruskal-Wallis test followed by Dunn’s multiple comparisons test was used to calculate P-values. ns : P > 0.05; * : P ≤ 0.05; ** : P ≤ 0.01; *** : P ≤ 0.001; **** : P ≤ 0.0001. J: Box plots of distances of HSPH1 and HSPA1 RNA relative to SON, THRAP3 and DDX39B. n (RNA spots) = 168-257. RNPs from 5-8 nuclei used for this analysis. Kruskal-Wallis test followed by pairwise Wilcoxon test was used to calculate P-values. ns : P > 0.05; * : P ≤ 0.05; ** : P ≤ 0.01; *** : P ≤ 0.001; **** : P ≤ 0.0001. K: Box plots showing average fraction overlap (Manders coefficient) between RNA signals and THRAP3 or DDX39B foci / concentrations for HSPH1 and HSPA1 RNAs. White squares represent mean and central line represents median. (n (number of nuclei): HSPH1 18; HSPA1 18). Kruskal-Wallis test followed by pairwise Wilcoxon test was used to calculate P-values. ns : P > 0.05; * : P ≤ 0.05; ** : P ≤ 0.01; *** : P ≤ 0.001; **** : P ≤ 0.0001. (A-D) Scale bar=1 μm. (E-H) Scale bar= 5 μm main image; 1 μm inset. (A-H) Regions highlighted within white boxes are shown as insets on the right.\

Overall, these results from both transgene arrays and endogenous genes indicate that the transcripts from intron-containing genes preferentially associate with PSTP within the interchromatin space while transcripts from both intron-containing and intronless genes show close association with PSDP.

### THRAP3 and DDX39B networks are largely maintained after NS knockdown

We next asked whether the maintenance of PSTP and PSDP networks depends on NS. One possibility is that NS nucleate these PSTP and PSDP networks and therefore these perispeckle networks would require NS for maintenance. Alternatively, these perispeckle networks might be maintained independently of NS, possibly even with NS then preferentially nucleating within regions of high local PSDP protein concentrations.

Previous work had concluded that SON and SRRM2 are core NS proteins and essential for NS formation (Ilik et al., 2020). We used simultaneous degradation of both SON and SRRM2 using an improved auxin-inducible degron (AID2) system (Fig. 8A, SFig. 8A) (Yesbolatova et al., 2020). Both endogenous alleles of SON and SRRM2 in HCT116 cells were modified with degron tags and mClover and mCherry fluorescent tags, respectively. Fixed and live-cell microscopy both show NS, identified by SON and SRRM2 concentrations, shrinking and then in most cases disappearing within 0-4 hours of analog 5-Ph-IAA treatment (Fig. 8B-C, SFig. 8B). Live-cell imaging reveals NS initially shrinking, while maintaining elevated concentrations of SON during this shrinkage (Fig. 8B, 20-40 mins), consistent with SON existing as a condensate within NS.

**Figure 8:**
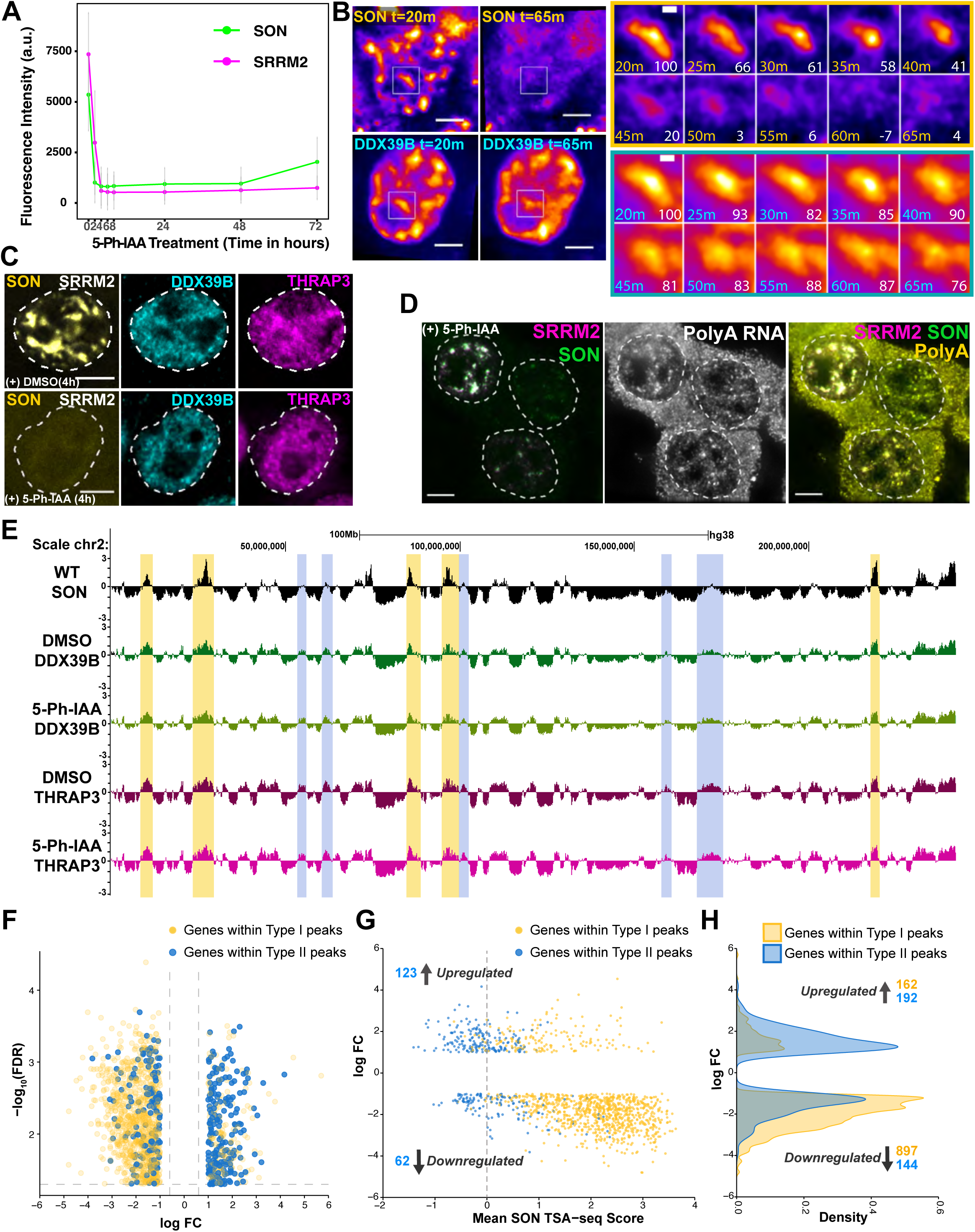
Perispeckle networks and genome positioning are largely maintained after NS depletion but NS-proximal genes are downregulated. A: SON (mClover, green) and SRRM2 (mCherry, magenta) intensities (y-axis) as a function of time (x-axis, hours) after 5-Ph-IAA treatment as measured by flow cytometry reveals maximum double knockdown after ∼ 4 hours in these HCT116 cells. B: Live-cell imaging reveals DDX39B distribution shows minimal changes during process of NS elimination (Video 6). Left: Pseudocolor intensity mapping of mClover-SON (endogenous gene expression, top panels) versus HaloTag-DDX39B (transiently expressed, bottom panels) at 20 (left) versus 65 (right) minutes after 5-Ph-IAA induced double knockdown (DKD) of SON and SRRM2. Right: Inset frames show SON (top) or DDX39B (bottom) over a single NS region at different times (minutes, m) after DKD. Percentages of SON (top) or DDX39B (bottom) remaining over NS region relative to the first frame intensities are shown as white numbers (bottom right, each frame). Scale bar-5 μm main image; 1 μm inset. C: Immunostaining of DDX39B (cyan) and THRAP3 (magenta) without (top panel) or with 5-Ph-IAA treatment for 4 hours (bottom panel). SON (yellow) and SRRM2 (gray) marked NS undergo degradation within 4 hours. D: Loss of nuclear poly-A RNA concentrations within NS occurs after NS elimination by DKD (5-Ph-IAA treatment). Nucleus (upper left) with significant SON (green) and SRRM2 (magenta) still remaining in NS also still shows poly-A (middle panel, grey; right panel, yellow) FISH signal accumulations within NS; dispersal of poly-A accumulation is seen in two nuclei (right) with greater SON and SRRM2 DKD. E: DDX39B (green) and THRAP3 (magenta) TSA-seq profiles of chromosome 2 in HCT116 cells engineered with AID2-inducible degron tagged SON and SRRM2 alleles after DMSO (dark colors) versus 5-Ph-IAA (light colors) treatment for 4 hours. SON TSA-seq (black) from WT cells is shown in comparison. Type I (yellow highlights) and Type II (blue highlights) peaks show little or no change in DDX39B and THRAP3 peak amplitudes after DKD. F-H. Differentially expressed genes (DEG) near SON TSA-seq Type 1 peaks show a strong bias towards downregulation. In contrast, DEG near SON TSA-seq Type 2 peaks show bias towards upregulation, particularly for SON TSA-seq peak amplitudes less than or equal to zero. F: Volcano plot showing log2 fold expression changes versus log10 (False Discovery Rate) for DEG present +/- 500 kb from SON TSA-seq Type I (yellow) and Type II (blue) peaks in HCT116 cells after 5-Ph-IAA-induced SON-SRRM2 degradation. G: Scatterplot showing mean SON TSA-seq score versus log2 fold changes for DEG present +/- 500 kb of Type I (yellow) and Type II (blue) peaks. Numbers of DEG flanking Type II peaks with SON TSA-seq peak amplitudes less than or equal to zero (left of grey dashed vertical line) are indicated in blue. H: Density plot with log2 fold change of upregulated and downregulated genes within Type I and Type II regions. Type I: 162 upregulated genes and 897 downregulated genes; Type II: 192 upregulated genes and 144 downregulated genes.

Immunostaining of DDX39B and THRAP3 shows similar staining networks both before and after NS depletion (Fig. 8C). We used transfection to express a HaloTag-DDX39B construct and performed live-cell imaging after two weeks of drug selection.

The cells were imaged 20-25 minutes after adding 5-Ph-IAA-containing media. These live-cell movies reveal a ∼20% reduction of DDX39B intensity within the regions formerly occupied by NS after SON and SRRM2 double knockdown (DKD); a concomitant increase in DDX39B intensities is seen in the network staining outside of the regions formerly occupied by NS (Fig. 8B).

At later time points (>45 mins), NS SON levels drop to just a few percent of their value at 20 mins (Fig. 8B), while DDX39B levels within the same NS decrease by only ∼25%. Importantly, the overall nuclear DDX39B intranuclear distribution is largely maintained, including the DDX39B concentrations over regions previously occupied by SON in NS.

The previous conclusion that SON and SRRM2 are required for NS nucleation and maintenance was based on the dispersion of several other NS markers throughout the nucleus after simultaneous SON and SRRM2 DKD using siRNA (Ilik et al., 2020; Zhang et al., 2024). An alternative explanation, however, could be that SON and SRRM2 are required only for recruitment of this subset of NS components but not for NS nucleation and maintenance per se. Our observation of an initial shrinking of the NS after SON and SRRM DKD using AID2 (Fig. 8A, B), simultaneous with retention of elevated SON and SRRM2 concentrations within these shrinking NS, is consistent with the shrinkage and then disappearance of an actual condensate dependent on SON and SRRM2, thus supporting a primary role of SON and SRRM2 in nucleating and maintaining NS.

NS have been operationally defined as the light microscopy equivalent of the Interchromatin Granule Clusters (IGCs) first identified by electron microscopy; these granules were demonstrated by immunostaining to contain RNA (reviewed in (Chen and Belmont, 2019)). At the light microscopy level, NS were also operationally defined by their local concentration of polyadenylated RNAs (Carter et al., 1991, 1993).

We therefore next examined the distribution of polyA-RNA after SON and SRRM2 acute DKD using RNA FISH. Indeed, we observed the dispersion of locally concentrated polyA-RNA “islands” after SON and SRRM2 DKD (Fig. 8D).

We therefore conclude first that NS are indeed depleted after SON and SRRM2 DKD and second that the DDX39B and THRAP3 perispeckle networks are largely maintained after NS depletion.

### Nuclear genome positioning relative to PSTP and PSDP is largely maintained independent of NS

Given the general alignment of THRAP3 and DDX39B with SON TSA-seq signals, plus the maintenance of PSTP and PSDP after NS elimination, we next asked whether chromosome positioning relative to PSTP and/or PSDP was dependent on maintenance of NS. We performed DDX39B and THRAP3 TSA-seq after 4 hours of 5-Ph-IAA induced SON and SRRM2 DKD, the earliest time point we had sampled showing maximum KD of both SON and SRRM2 expression (Fig. 8A). We observed little change in chromosome positioning relative to PSTP and PSDP after this several hour loss of SON and SRRM2 and NS, including over gene expression hot-zone chromosome regions corresponding to both Type I and II SON TSA-seq peaks (Fig. 8E, SFig. 8B-E).

Overall, both the THRAP3 and DDX39B TSA-seq reveal little change after NS elimination. Quantitative comparisons show a slight, largely statistically insignificant increase in DDX39B TSA-seq values after NS elimination (SFig. 8B). Conversely, THRAP3 TSA-seq shows a small overall decrease in signal after NS elimination, which does score as statistically significant (SFig. 8C); this small change in THRAP3 TSA-seq scores appears to reflect a systematic variation in scaling across the genome after SON and SRRM2 DKD. Specifically, THRAP3 TSA-seq peaks are decreased in amplitude slightly while valleys are deepened slightly. The reasons for these subtle genome-wide changes in DDX39B and THRAP3 TSA-seq score amplitudes remain unclear but may be related to small shifts in DDX39B and THRAP3 intranuclear distribution after NS KD.

Despite these small changes in scaling of the DDX39B and THRAP3 TSA-seq signals, we do not see any significant or specific changes in the overall patterns of peaks corresponding to NS and perispeckle pattern associated domains (Fig.8E, SFig.8B-E).

Finally, we asked how genes differentially positioned relative to NS might change their expression as a function of SON and SRRM2 DKD. SON depletion is known to arrest cell growth through perturbation of the cell cycle, particularly during mitosis (Sharma et al., 2010; Ahn et al., 2011). To minimize changes in gene expression due to changes in cell cycle progression, we combined an extended 16 hr DKD of SON and SRRM2 with a cell-cycle arrest in late G1/early S-phase. We applied the DKD in cells as they progressed from G1-arrest into a late G1/early S phase block (see Methods).

RNA-seq analysis revealed that NS-proximal genes within +/- 500 kb of Type I SON TSA-seq peaks showed a systematic bias towards decreased gene expression after SON and SRRM2 DKD (Fig. 8F-H; Supplementary Table 2). In contrast, genes within +/- 500 kb of Type II SON TSA-seq peaks, showed no strong bias towards decreased versus increased differential gene expression after SON and SRRM2 DKD (Fig. 8F-H; Supplementary Table 2). We note that the SON TSA-seq peak-calling used Hi-C subcompartment identity to classify Type I versus Type II peaks, which caused some degree of misclassification of SON TSA-seq positive peaks (Type I) versus “peak-within-valley” SON TSA-seq local maxima (Type II). Instead, defining Type II peaks as those with SON TSA-seq scores ≤ 0.0 (Fig. 8G, dashed line), we observe a more pronounced ∼2-fold bias of increased (123 genes) versus decreased (62 genes) differentially expressed genes within Type II peaks after NS KD (Fig. 8G, left of dashed vertical line).

Thus, we conclude that chromosome positioning relative to PSTP and PSDP networks, including regions originally near NS defined by SON TSA-seq peaks, remain unchanged for at least several hours after NS elimination. This observation raises the possibility that the perispeckle patterns might maintain genome organization independent of the NS defined by SON and SRRM2. Despite this maintenance of genome organization of Type I and II gene expression hot-zones, genes in Type I hot-zones do appear dependent on NS to sustain their normal levels of gene expression.

## Discussion

### Summary

We set out to resolve an apparent paradox. On the one hand, live-cell imaging (Kim et al., 2020) and RNA FISH (Zhang et al., 2021; Alexander et al., 2021) both suggested actual contact with NS was required for gene expression amplification. On the other hand, a strong correlation was observed genome-wide between changes in gene expression and positioning of these genes relative to nuclear speckles, even when these changes in positioning did not involve changes in actual nuclear speckle close association (Zhang et al., 2021; Gholamalamdari et al., 2025). To reconcile these two distinct observations, we hypothesized the existence of a previously unidentified gene expression hub(s) that was itself spatially correlated with but distinct from NS.

Here we describe the identification of two perispeckle patterns, characterized by the concentrations of specific proteins, which are both tightly spatially correlated with NS (Fig. 9). PSTP components surround NS, defined by SON and SRRM2 immunostaining, and in some cases may colocalize but not concentrate within NS, while PSDP components concentrate within NS. More importantly, both extend outwards and throughout the nuclear, DNA-depleted interior in thread-like patterns from NS. Together PSTP and PSDP subdivide the ICS. Meanwhile, other proteins that concentrate within the ICS overlap with neither PSTP nor PSDP, suggesting that these two perispeckle patterns together still occupy only a fraction of the total ICS; this would leave substantial additional space which other condensates might occupy.

**Figure 9:**
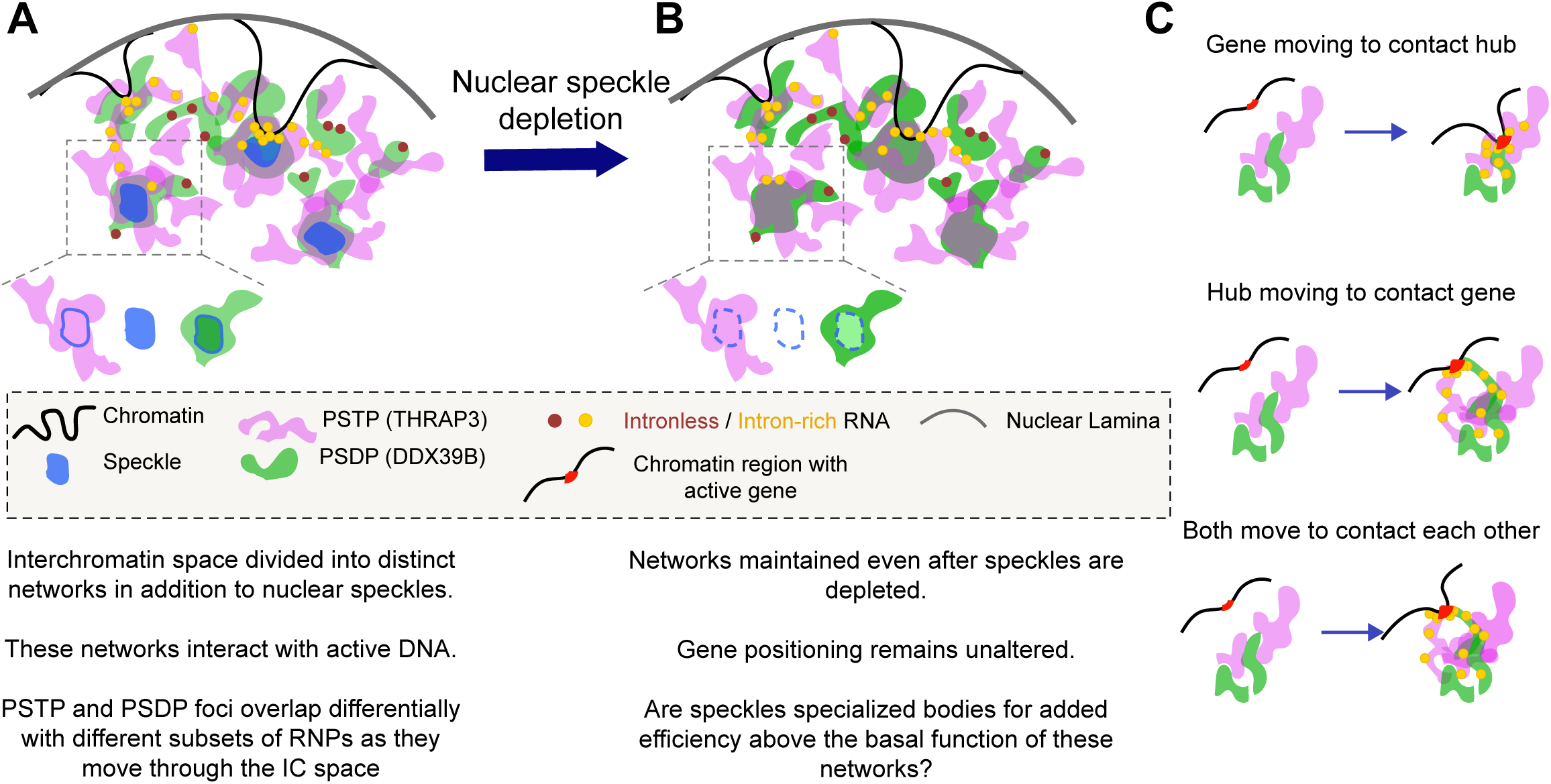
Model showing organization of genes relative to NS and DDX39B (PSDP) and THRAP3 (PDTP) perispeckle patterns. A: We propose NS versus perispeckle patterns correspond to two different nuclear niches acting as gene expression “hubs” for two different sets of genes. These sets of genes are contained within chromosomal regions with similarly elevated levels of gene expression but differential localization relative to NS. Genes near SON TSA-seq Type I peaks (Type I gene expression hot-zones) associate closely with NS (blue). Genes near SON TSA-seq Type II peaks (Type II gene expression hot-zones) mostly locate away from NS but near perispeckle patterns marked by THRAP3 (magenta, PSTP) and DDX39B (green, PSDP). Transcripts from intron-rich genes (yellow) associate with PSTP and PSDP while transcripts from intronless genes (red) associate only with PSDP. B: After NS depletion, genome organization relative to PSTP and PSDP remains largely the same, although many of the Type I set of genes are downregulated in the absence of NS, suggesting that NS confer added gene expression amplification for this set of genes. C: We imagine that positive regulation of gene expression may result from one set of genes associating with NS and the other set of genes associating with perispeckle patterns. Previous observations of gene movements from the nuclear lamina towards the nuclear interior and/or towards NS would suggest this occurs, at least in some cases, through genes moving towards either of these two hubs (top), but this might also occur through NS and/or perispeckle patterns moving or extending towards genes (middle), nucleating near genes (not shown), or a combination of gene and hub movements towards each other (bottom).

PSTP and PSDP appear to represent an alternate gene expression hub in the sense that a large fraction of the chromosomal regions with high gene expression levels not localized near NS are instead localized very close to the thread-like patterns of PSTP and PSDP. Similarly to nuclear speckles, PSTP and PSDP contain a mixture of components involved in various steps of gene expression, especially RNA splicing and export. Additionally, we show the preferential association of RNAs from two model intron-containing genes with PSTP as compared to the RNAs from two model intron-less genes. In contrast, the RNAs from both intron-containing and intronless genes are closely associated with PSDP. Thus, we speculate that the PSTP and PSDP perispeckle networks may additionally facilitate a spatial coupling between the machinery of transcription and RNA processing and export for a subset of active genes and their transcripts (Fig. 9).

### Possible resolution of our paradox

We propose that the close association of highly active chromosomal regions not near NS with these perispeckle patterns may resolve the apparent paradox between our previous live-cell imaging and genomic analysis of the relationship of active genes relative to NS. Because of the close spatial arrangement of PSTP and PSDP relative to NS, repositioning of chromosomal regions with gene activation to bring them into contact with these perispeckle patterns (Fig. 9, right panel) will also bring these active chromosomal regions closer to NS on average over the cell population. Most RNA Pol II Ser2p and Ser5p foci localize within several hundred nm of PSDP and, especially, PSTP foci, even those positioned away from NS. Similarly, chromosomal gene expression hot-zones, identified as Type I and Type II SON TSA-seq peaks, localize within several hundred nm of PSTP and PSDP foci. This is true even when positioned away from NS, as is the case for the majority of alleles of Type II SON TSA-seq peak regions. Moreover, as predicted in our original model (Fig. 1A), we showed that a chromosome gene expression hot-zone corresponding to a Type II SON TSA-seq was the closest chromosome region to a PSTP or PSDP foci over the entire chromosome iLAD trajectory stretching from the nearest LAD to this Type II SON TSA-seq peak.

### Perispeckle patterns are dynamic networks rather than stable nuclear bodies

In our original hypothesis, we had envisioned Type II SON TSA-seq peaks as interacting with an alternative nuclear body/compartment of comparable stability to NS. Instead, we observe both PSTP and PSDP networks away from NS to be highly dynamic over a time period of seconds to tens of seconds, even though we see their repeated recurrence in similar spatial patterns on a time scale of minutes such that time-averaging reveals a more definite network pattern. In a companion paper, we more thoroughly examine dynamics of PSDP components DDX39B and MAGOH and show that their preferential concentration into stable time-averaged patterns disappears with ATP-depletion and at least partially with transcriptional inhibition (Kim et al., 2025). We also show directional, bulk transfer of SON condensates between NS within PSDB network connections (Kim et al., 2025).

### Relationship of DDX39B perispeckle pattern to NS

Despite the close spatial relationship of PSTP and PSDP to NS, both THRAP3 and DDX39B distributions are largely maintained after NS disruption. SON and SRRM2 were considered to be required for NS integrity because their double KD (DKD) led to dispersal of several other known NS proteins (Ilik et al., 2020). DDX39B, and possibly other PSDP components, retain the same high concentrations within bodies that originally colocalized with NS even after SON and SRRM2 KD that is reported to eliminate NS (Ilik et al., 2020). Should these high-concentration DDX39B bodies then be considered as NS that persist after SON and SRRM2 DKD with SON and SRRM2 necessary only for retention of a subset of NS proteins?

NS were described originally as clusters of RNPs and foci of polyA accumulation. From this functional view of NS as concentrators of RNPs, we therefore suggest that the observed loss of NS-associated poly-A RNA accumulations after SON and SRRM2 DKD means NS are indeed eliminated after SON and SRRM2 KD. Just as nucleolar GC proteins such as nucleolin can mix with the FC (Lafontaine et al., 2021), we suggest PSDP components have the ability to mix with NS components. Similar to how the dense fibrillar component and fibrillar centers (FC) form within the granular compartment (GC) of nucleoli, we suggest that NS form within accumulations of PSDP components

A second question is whether gene positioning is controlled not by NS per se but by PSDP, given that gene positioning is largely maintained for at least several hours after NS disruption, and what functions NS may impart separate from PSDP. The observation of gene expression amplification with NS contact (Kim et al., 2020; Alexander et al., 2021; Chaturvedi and Belmont, 2024), plus the observed reduced expression specifically of a subset of genes positioned very close to NS after SON and SRRM2 KD, suggests that NS are indeed required for the increased expression of a subset of NS proximal genes.

### Connections of perispeckle patterns to RNA processing and trafficking

Similar to NS, which are enriched especially for proteins related to multiple levels of RNA processing, modification, and export, the perispeckle pattern marker proteins we identified here are all related to RNA splicing and export. Our demonstration of close association of RNA with these two perispeckle patterns raises the interesting possibility that these perispeckle patterns might be pathways for mRNAs moving away from the gene and trafficking through the nucleus.

Among the PSTP marker proteins, THRAP3 (TRAP150) was described as a member of the TRAP/Mediator complex (Ito et al., 1999) but also associates with mRNA export factor TAP (Lee et al., 2010) and has been shown to play roles in pre-mRNA splicing (Lee et al., 2010; Lee and Tarn, 2014; Vohhodina et al., 2017), mRNA degradation (Lee et al., 2010), and R-loop resolution (Kang et al., 2021). BCLAF1 has known roles in pre-mRNA splicing (Vohhodina et al., 2017), nuclear retention of RNAs (Varia et al., 2013) and as a transcriptional repressor (Shang et al., 2022) . PLRG1 is a component of the cell division cycle 4-like (CDC5L) complex which is part of the spliceosome (Ajuh, 2000) and is involved with alternative splice-site selection (Llères et al., 2010), as well as DNA repair (Zhang et al., 2005) and the ATR-response (Zhang et al., 2009). During development, PLRG1 regulates proliferation and apoptosis (Kleinridders et al., 2009). Interestingly, we observed the mRNAs from intron-rich genes, but not intronless genes, associating closely with PSTP.

Among the PSDP marker proteins, DDX39B is a DEAD-box RNA helicase already known to regulate release of spliced mRNAs from NS (Hondele et al., 2019; Taniguchi and Ohno, 2008; Luo et al., 2001; Fan et al., 2018). DDX39B is also essential for R-loop resolution (Pérez-Calero et al., 2020) and is a member of the TREX export complex (Dias et al., 2010; Wang et al., 2018; Shen et al., 2008). RBM8A (Y14) and MAGOH are both members of the core Exon Junction Complex (EJC) that assembles on pre-mRNAs over the junction of two exons that are spliced together (Le Hir, et al. 2000) and provides binding platform for additional factors involved in mRNA export and nonsense-mediated decay (NMD) (Le Hir, Hervé, et al. 2001).

Interestingly, DDX39B previously was suggested to regulate the entry and release of RNAs from phase-separated condensates in an ATP-dependent manner (Hondele et al., 2019). Immunogold EM against DDX39B coupled to EDTA-regressive staining showed DDX39B associating at the periphery of IGCs and thread-like, likely RNA-containing projections extending from IGCs into the interchromatin space (Kota et al., 2008). DDX39B is evolutionarily-conserved and is essential for the export of spliced RNA and intronless RNA (Jensen et al., 2001; Gatfield et al., 2001; Kota et al., 2008; Dias et al., 2010; Wang et al., 2018). Higher-order assembly of export-competent mRNPs is also mediated by DDX39B (Xie et al., 2023). Cosistent with a role of DDX39B in nuclear RNA export, we observed mRNAs from both intron-rich and intron-less genes associating closely with PSDP.

All together, these observations raise the hypothesis of RNPs being moved sequentially through different spatially organized “processing” condensates before and during export out of the nucleus.

## Future Directions

Our description of perispeckle patterns suggest multiple avenues for future investigation. Of central interest will be going beyond correlation to test causality between chromosomal and RNA association with PSTP and PSDP networks with modulation of gene expression, including RNA export. Future live-cell imaging will be valuable to test the temporal correlation between contact with PSTP and/or PSDP and changes in gene expression and/or RNA movements as well as monitoring changes in gene expression after acute knockdown of PSTP and PSDP components. Beyond gene expression, we anticipate future investigations into the possible role of perispeckle patterns in DNA repair and genome stability given that multiple PSTP and PSDP components have been implicated in both R-loop resolution (THRAP3, DDX39B, POLDIP3) (Kang et al., 2021; Pérez-Calero et al., 2020; Björkman et al., 2020) and DNA repair and genome stability (THRAP3, BCLAF1, RBM8A, DDX39B, PLRG1) (Vohhodina et al., 2017; Yu et al., 2023a; Chuang et al., 2019; Xu et al., 2020; Choi et al., 2023).

Two additional functions of perispeckle patterns are suggested by our work and others. First, a role of PSDP in bulk trafficking of NS components is suggested by a companion study. Using live-cell imaging, we demonstrated the directional bulk transfer of SON protein as small condensates between NS in ATP-dependent “connections” marked by dynamic DDX39B, MAGOH, and SON multi-phase protein concentrations whose interactions form a more stable unit that any of the individual components (Kim et al., 2025). Second, perispeckle patterns may play a role in genome stability, as multiple components of both PSTP and PSDP have been implicated in both R-loop resolution (THRAP3, DDX39B, POLDIP3) (Kang et al., 2021; Pérez-Calero et al., 2020; Björkman et al., 2020) and DNA repair and genome stability (THRAP3, BCLAF1, RBM8A, DDX39B, PLRG1) (Vohhodina et al., 2017; Yu et al., 2023a; Chuang et al., 2019; Xu et al., 2020; Choi et al., 2023).

Finally, although we focused here on PSTP and PSDP, our initial screening suggested that there will be additional perispeckle patterns/ protein condensates that further subdivide the interchromatin space. We therefore suggest additional investigations will be needed to fully appreciate the integration of these dynamic protein condensates with each other and with important nuclear functions.

## Methods

### Cell lines and cell culture

U2OS and HCT116 cells were obtained through the ATCC, expanded through pyramid stocking, and maintained as per the 4D Nucleome SOP (https://data.4dnucleome.org/resources/experimental-resources/cell-lines). U2OS cell lines containing E6 β-globin and mini-Dystrophin A1 transgenes were described previously (Mor et al., 2010; Brody et al., 2011).

All cells were grown at 37°C in a humidified incubator with 5% CO2. Specifically, both U2OS and HCT116 cells were cultured in McCoy’s 5A medium (UIUC cell media facility) supplemented with 10% fetal bovine serum (FBS) (VWR, cat# 97068-091, Lot #035B15). Cells were passaged (or harvested) at 70%-80% confluency using Trypsin-EDTA (0.05%) (Thermo Fisher Scientific, cat# 25300062).

U2OS cell lines with the E6 β-globin and mini-Dystrophin transgenes were cultured in low glucose containing DMEM (UIUC cell media facility). The E6 β-globin cell line was maintained with 10 μg/ml Puromycin (Puromycin Dihydrochloride, Gibco, cat# A1113803) and 0.54 mg/ml Zeocin (Gibco, cat# R25001). The mini-dystrophin A1 cell line was cultured with 50 μg/ml Hygromycin (Gibco, cat#10687010) and 0.54 mg/ml Zeocin (Gibco, cat# R25001). Cells were passaged using Trypsin-EDTA (0.05%) (Thermo Fisher Scientific, cat# 25300062).

For all cell lines, cells were cultured for a maximum of 10-12 passages after thawing.

### cDNA Constructs and screening

For the screening of selected protein candidates, PCR amplicons were prepared from cDNA. For genes with multiple transcript variants, the longest protein-coding transcript sequence was chosen for PCR amplification and cloned into pEGFP-C1 vector as described previously (Dopie et al., 2020). Briefly, total RNA was isolated using the RNeasy kit (Qiagen, cat#74104) and 1μg total RNA was reverse transcribed using the Protoscript First Strand cDNA Synthesis Kit (New England Biolabs, cat# M0368S) to generate cDNA. Using primers containing the appropriate restriction enzyme sites (Supplementary Table 1), cDNA amplicons were PCR amplified with Q5 High-Fidelity DNA Polymerase (New England Biolabs, cat# M0491S), purified and cloned into pEGFP-C1 (originally obtained from Clonetech). The sequences of all plasmids generated were validated by Sanger sequencing before their use.

For transfections, ∼120,000 U2OS cells were seeded per well in a 6-well plate containing 12mm diameter #1.5 coverslips (Electron Microscopy Sciences, cat#72230-01) and the next day, 500 ng of plasmid was transfected using 5ul Lipofectamine 2000 (Thermo Fisher Scientific, cat#11668019) in OptiMEM (Thermo Fisher Scientific, cat# 31985070) as per the manufacturer’s instructions. After 24 hours, cells were fixed with freshly prepared 4% paraformaldehyde (Sigma, cat#P6148) in calcium and magnesium free phosphate buffered saline (CMF-PBS) for 10 mins at room temperature (RT). Cells were washed for 5 mins 3 times with CMF-PBS and then either mounted directly using Mowiol-DABCO antifade mounting medium or processed further.

### Antibody generation

A rabbit polyclonal anti-DDX39B antibody (Pacific Immunology) was generated against the 24 amino acid synthesized peptide (Cys)-DRFEVNISELPDEIDISSYIEQTR, conserved across human, mouse and Chinese hamster DDX39B protein sequences.

Briefly, two rabbits were immunized once with AdjuLiteTM Complete Freund’s Adjuvant followed by three rounds of AdjuLiteTM Incomplete Freund’s Adjuvant immunizations. The first production bleed was validated by ELISA against the peptide used for immunization. After validation, serum from both rabbits was pooled prior to affinity purification. 25 mL pooled serum was subjected to affinity purification using the same target peptide. This affinity-purified antibody was diluted in 50% glycerol for long-term storage at a concentration of 0.6 mg/ml. The antibody was further validated by demonstrating its colocalization after immunostaining with the mCherry-DDX39B signal in a U2OS cell line with the endogenous DDX39B gene tagged with mCherry (See “Cell Line Construction” Section, below).

### Immunostaining

Cells were fixed with freshly prepared 4% (w/v) paraformaldehyde solution in CMF-PBS (pH 7.0) (Sigma-Aldrich, cat# P6148) for 10 mins at RT. Coverslips were washed three times with CMF-PBS and cells were permeabilized with 0.5 % Triton X-100 in CMF-PBS for 10 mins at RT. Washes for 5 mins with CMF-PBS were repeated three times followed by blocking for 1 hr at RT in 5% (v/v) goat serum solution prepared in 0.1 % Triton X-100 in CMF-PBS. Primary antibodies were diluted as stated (Supplemetary Table 1) in blocking buffer and applied to coverslips overnight at 4°C for 12-16 hours in a humidified chamber. Coverslips then were washed for 5 mins three times using 0.1 % Triton X-100 in CMF-PBS at RT. Secondary antibodies were diluted as stated (Supplementary Table 1) in blocking buffer and applied to coverslips for 1 hr at RT in a humidified chamber before 5 min washes with 0.1 % Triton X-100 in CMF-PBS repeated three times. Coverslips were mounted using Mowiol-DABCO antifade mounting medium without or with DAPI (Sigma-Aldrich cat#D9542) (10ug/ml).

### Cell line construction

Knock-in cell lines containing endogenous gene sequences tagged with different fluorescent proteins were constructed using CRISPR/Cas9 as described previously (Gholamalamdari et al., 2025). For gRNA synthesis, a 2-step overlap PCR strategy was employed. In the first step, a forward primer including the T7 promoter sequence followed by the target specific sequence 20bp immediately upstream of the PAM sequence and a reverse primer containing the tracrRNA template sequence (5’-AAAAGCACCGACTCGGTGCCACTTTTTCAAGTTGATAACGGACTAGCCTTATTTTAA CTTGCTATTTCTAGCTCTAAAAC -3’) (Supplementary Table 1) were used. This first step generated a 120bp T7-gRNA-tracr fragment. In the second step, we amplified this DNA template using a T7 promoter sequence forward primer (5’-TAATACGACTCACTATAG-3’) and a reverse primer sequence (5’-AAAAGCACCGACTCG-3’).

This DNA template was gel purified (QIAGEN # 28704) and used for an in vitro-transcription reaction to generate gRNA. 1 μg of DNA template was incubated with 2.5mM ribonucleotides (NTP, New England Biolabs cat#N0466S), 1U/μl Ribolock RNase inhibitor (Thermo Fisher Scientific, cat# EO0381), 8 μl T7 RNA Polymerase (purified in the laboratory) in 50 μl reaction volume in IVT buffer (50mM Tris-HCl, pH 7.6, 15mM magnesium chloride, 5mM dithiothreitol (DTT) and 2mM spermidine). The reaction was incubated at 37°C for 6-16 hours in a thermocycler. In-vitro transcribed product was treated with 2.5 U DNase I (2 U/μl; New England Biolabs cat# M0303S) at 37°C for 15 mins to remove the DNA template prior to purification. gRNA was purified using the RNeasy plus kit (QIAGEN cat# 74134). Purified gRNAs eluted in RNase-free water (Accugene cat#51200) were stored as aliquots at -80°C.

2 μg of target-specific gRNAs were complexed with 3μg purified Cas9 Nuclease V3 (Integrated DNA Technologies, cat# 1081058). gRNA sequences and primers used for homology arms of Donor DNA sequences are provided in Supplementary Table 1. Primers 70-90 bp long with their 5’ end modified with an amine group coupled to a C6 linker (Yu et al., 2020) were used for generating donor DNA with homology arms including sequences upstream and downstream of the target genomic DNA cut site; (Supplementary Table 1). PCR amplification of the donor DNA used Q5 High-Fidelity DNA Polymerase (New England Biolabs, cat# M0491S) and was applied to a plasmid or BAC template DNA containing sequences coding for the fluorescent protein and linker peptide to be added to the NH2-terminus of the target protein (Supplementary Table1). This donor DNA was purified after the PCR using magnetic beads (MAGBIO, HighPrep PCR, cat#AC-60050). Nucleofection was performed using an Amaxa Nucleofector II device (Lonza) as per the manufacturer’s instructions.

After 5 days of recovery, fluorescence-positive cells were sorted using the FACS ARIA II sorter at the UIUC Roy J. Carver Biotechnology Center. Using this approach, the “U2OS EGFP-SON C5” cell clone was first generated. U2OS EGFP SON C5 was then further modified to generate the “U2OS EGFP-SON C5 mCherry-DDX39B C11” cell clone and the “U2OS EGFP-SON C5 mCherry-THRAP3 gRNA#1” mixed clonal cell line. Similarly, a U2OS HaloTag-SON” mixed clonal cell line was generated and then further modified to generate the “U2OS HaloTag-SON EGFP-THRAP3 gRNA#1” mixed clonal cell line (Supplementary Table1).

HCT116 SON and SRRM2 double-degron cell line (expressing mAID-Clover-SON and SRRM2-mAID-mCherry) was generated by CRISPR-Cas9-mediated genome editing as previously described (Yesbolatova et al. Nat. Commun., 2020; Saito et al. Current Protocols, 2021). Briefly, we initially fused mAID-mCherry to the C-terminus of SRRM2. For this, we designed a CRISPR-Cas9 plasmid for targeting the SRRM2 locus (5’-AGGTCTCCATAAATTGTCTT(TGG)-3’). A donor plasmid harboring mAID-mCherry and a selection marker (BSD) with two homology arms (approximately 500 bp each) was constructed. HCT116 cells stably expressing OsTIR1(F74G) from the AAVS1 locus were transfected with the CRISPR and donor plasmids. Clones surviving blasticidin selection were screened for biallelic by genomic PCR, and expression of the fusion protein was confirmed by immunoblotting. Subsequently, we fused mAID-Clover to the N-terminus of SON. For this, we designed a CRISPR-Cas9 plasmid for targeting the SON locus (5’-AGAGAACGGAGCGGACGCCA(TGG)-3’). A donor plasmid harboring HygroR-P2A-mAID-mClover with two homology arms was constructed. The SRRM2-mAID-mCherry cells were transfected with the CRISPR and donor plasmids for tagging SON. Clones surviving hygromycin selection were screened for biallelic insertion of the tagging construct by genomic PCR, and expression of the fusion protein was confirmed by immunoblotting. To induce degradation of SON and SRRM2, 5-Ph-IAA was added to the final concentration of 1 µM.

### Transcriptional inhibition treatment

30,000 U2OS cells were seeded per #1.5 12 mm diameter coverslip (Electron Microscopy Sciences, cat# 72230-01) in a 24-well plate. After two days, cells were treated with 50 µg/ml DRB (Sigma-Aldrich, cat# D1916), 50 µg/ml α-amanitin (Sigma-Aldrich, cat# A2263), or 0.1 µg/ml triptolide (Sigma-Aldrich, cat# T3652) added in McCoy’s 5A media for the time indicated and fixed. For heat shock experiments, cells were seeded on #1.5 12 mm coverslips placed within 35mm dishes (Corning, cat# 430165) at 120,000 cells per dish. After two days, cells were incubated in a water bath at 42°C after sealing the dishes with parafilm (Parafilm, cat# S25929) for the indicated time and fixed immediately.

### Tyramide Signal Amplification-sequencing (TSA-seq)

A detailed protocol for TSA-seq is described elsewhere (Zhang et al., 2022). For high-resolution mapping, we combined the use in the original TSA-seq 1.0 procedure (Chen et al., 2018) of both sucrose and Dithiotheitol (DTT) to increase spatial resolution with the higher tyramide concentration and longer TSA reaction time used in the TSA-seq 2.0 protocol, which saturated the protein TSA labeling but increased the DNA tyramide labeling without saturation (Zhang et al., 2021). More specifically, we used a TSA reaction buffer prepared in CMF-PBS with 50% sucrose: 1.5 mM DTT (Millipore Sigma, cat#233155-10GM), and 0.33 mM tyramide-biotin (Apex-Bio, cat# A8011) with a reaction time of 30 mins. By boosting the tyramide concentration beyond previous TSA-seq 2.0 TSA reaction conditions, we compensated for the decreased TSA-labeling caused by the addition of the DTT, while keeping the nucleic acid labeling undersaturated as judged by microscopy as described previously (Zhang et al., 2021).

For each TSA-seq sample in U2OS cells, cells were grown in two T-300 flasks (∼15 million cells per T-300 flask). For each TSA-seq sample in HCT116 cells, cells were grown in one T-150 flask (∼30-40 million cells). For the TSA-seq of the HCT116 mAID-Clover-SON SRRM2-mAID-mCherry CMV-OsTIR1(F74G) cell line, cells in T-150 flasks were first treated with either DMSO (Avantor # IC0219605590) or 5μM 5-Ph-IAA (Tocris Biosciences #7392) for 4 hours before proceeding to the TSA-seq procedure.

All TSA-seq libraries were prepared using 3-10 ng of purified DNA using the TruSeq ChIP Library Preparation Kit (Illumina, cat# IP-202-1012) as per the manufacturer’s instructions. Libraries were PCR-amplified for 8 cycles and purified using AMPure XP magnetic beads (Beckman Coulter, cat# A63881). Concentrations of individual libraries were estimated by qPCR and then pooled accordingly. They were then sequenced from one end of the fragments on a NovaSeq X Plus with V1.0 sequencing kits. FASTQ files were generated and demultiplexed with the bcl-convert v4.1.7 Conversion Software (Illumina). TSA-seq normalization, scores and identification of changed regions were performed using a previously reported pipeline (Chen et al., 2018; Zhang et al., 2021).

### Oligolibrary design and probe synthesis

For DNA FISH of TSA-seq peaks, we used an oligolibrary containing ∼500 FISH probes targeted to DNA sequences within each ∼100 kb targeted peak region (Supplementary Table 1). Oligo probes (Supplementary Table 1) were designed using PAINTSHOP (https://paintshop.io/) as per published recommendations (Beliveau et al., 2012, 2017). Briefly, selected genomic region coordinates were uploaded and advanced probe settings specified as: no repeats allowed, max off-target score 15, max K-mer count 5, Min prob value 0 and probe set optimized to generate ∼500-523 probes per 100 kb target with “trim” setting for each region. Each probe set was appended with universal outer primers for each TSA-seq peak type, and a unique inner primer set for each 100 kb genomic peak region. The 3’ inner and outer primer was followed by the T7 promoter sequence.

For trajectory mapping, the distances between two consecutive ∼100 kb regions were kept at least 200 kb apart between the nearest edges of two neighboring regions. The outer primer was unique for “Apex” and “Arm” regions and each 100 kb genomic region had a unique inner primer set. The 3’ inner and outer primer was followed by the T7 promoter sequence. Otherwise, settings for probe selection were kept the same as for the TSA-seq peak probe selection.

For RNA-FISH, ∼100 probes were selected for MALAT1 RNA. The outer primer was unique for the target. The outer primer was followed by the T7 promoter sequence. Otherwise, settings for probe selection were kept the same as used for the DNA FISH probe selection.

The detailed protocol used for probe preparation is described elsewhere (Beliveau et al., 2017). Briefly, probes were amplified using PCR with Q5 High-Fidelity DNA Polymerase for 7-8 cycles, followed by in-vitro transcription using HiScribe T7 High Yield RNA Synthesis Kit (New England Biolabs cat# E2040S) and reverse-transcription using Maxima H Minus Reverse Transcriptase (Thermo Fisher Scientific cat# EP0751) as described previously (Beliveau et al., 2017).

2 μl of reactions were loaded on a urea-TBE gel to check for the desired ∼70-80% incorporation of reverse transcription RT primer (20bp) into the probes (∼150 bp). Probes were purified using Zymo DCC-25 columns as described previously (Beliveau et al., 2017).

2 μg purified probes were end-labelled using 12U Terminal Deoxynucleotidyl Transferase (TdT 20U/μl; Thermo Scientific, cat# EP0161) in TdT buffer (Thermo Scientific, cat# EP0161) with 108μM unlabelled dATP or dTTP and 54μM Biotin-dATP (Invitrogen #19524016) or Digoxigenin-11-dUTP (Roche #11093088910) respectively in a 25 μl reaction volume. Reactions were incubated at 37°C for 1hr in a thermocycler.

Probes were purified using the QIAquick Nucleotide Removal kit (QIAGEN, cat# 28306). Probes were eluted in 100 μL molecular biology grade water (Accugene, cat#51200) and human Cot-1DNA (Invitrogen #15279011) was added to each probe such that human Cot-1 DNA was 10x higher than the probe concentration. The resulting probe solutions were concentrated using a Vacufuge concentrator (Eppendorf) with the settings for aqueous solutions at 45°C for 45 mins until the probe solutions were reduced to ∼2-5 μl volumes. To these solutions, equal volumes of 100% formamide (Sigma-Aldrich, cat#F9037) and 2x hybridization buffer (20% w/v dextran sulphate (Sigma-Aldrich, cat# D8906) in 4x Saline-Sodium Citrate (SSC) buffer) are added such that the final probe concentrations are 25 ng/μl in hybridization buffer (10% w/v dextran sulphate in 2X SSC buffer with 50% formamide).

### DNA-FISH

Cells were fixed and immunostained as described previously (Solovei and Cremer, 2010). Briefly, cells were immunostained as described above with the exception of replacing the goat serum with 5% Bovine Serum Albumin (Sigma-Aldrich cat#A9418) for blocking and post-fixing after the immunostaining using 2% paraformaldehyde in CMF-PBS buffer. After three 5 min washes with CMF-PBS, coverslips were incubated with 20% glycerol (Fisher Scientific, cat# BP229-1) in CMF-PBS for 24 hours. Coverslips underwent 6 rounds of freezing and thawing in 20% glycerol in PBS with liquid nitrogen immersion followed by three 5 min washes with CMF-PBS containing 0.1% Triton X-100 (Thermo Fisher Scientific, cat# 28314).

Coverslips were then rinsed with 0.1 N HCl in molecular biology grade water (Accugene, cat#51200) and incubated in same at RT for 10 mins. Three 5 min washes with CMF-PBS containing 0.1% Triton X-100 were followed by immersion in 2x SSC containing 50% formamide at 4°C for at least 3 days and up to 15 days. For hybridization, oligolibrary probes (100-125ng of labelled oligoprobes in 10% dextran sulphate in 2X SSC buffer containing 50% formamide and 1μg of unlabelled human Cot-1 DNA (Invitrogen cat#15279011) were added to a glass slide and the 12 mm circular #1.5 mm coverslip was inverted and sealed using rubber cement (Elmers).

The slide was placed on a heating block at 75°C for 3 mins and then immediately moved to a humidified chamber at 37°C for 4 days of hybridization. Post-hybridization,coverslips were removed from the slide and washed for 10 mins three times with 2x SSC at 37°C followed by two 5 min washes with 0.1x SSC at 60°C.

Probes were detected using anti-Digoxin antibody (Jackson #200-602-156) or Streptavidin (Jackson #016-600-084) conjugated to Alexa 647 diluted 1:200 in 4% BSA in 4x SSC containing 0.2% Tween20 (Sigma-Aldrich, cat# P1379) at RT for 1 hr in a humidified chamber. Three 5 min washes were performed using 4x SSC with 0.2% Tween20 at 37°C. Coverslips were rinsed with CMF-PBS containing 0.1% Triton X-100 and mounted with DABCO containing mounting medium with/without DAPI.

### RNA-FISH

smRNA-FISH was performed as reported previously (Kim et al., 2020). All reagents used for this procedure were RNase free and/or diethyl pyrocarbonate (DEPC)-treated (Sigma-Aldrich cat# D5758). Cells were fixed with 4% paraformaldehyde for 10 mins at RT in CMF-PBS. Three 5 min washes in CMF-PBS were followed by permeabilization with 0.5% Triton X-100 in CMF-PBS for 10 min at RT followed by 3x 5 min washes in CMF-PBS. Cells were equilibrated using 10% formamide in 2X SSC for 30 min at RT.

smRNA FISH probes designed against HSPA1A (Kim et al., 2020), HSPH1 (Zhang et al., 2021), and oligolibrary probes designed against MALAT1 (Supplementary Table 1) were used at a final concentration of (∼300-500nM in hybridization buffer composed of 10% formamide, 10% (w/v) dextran sulphate (Sigma-Aldrich, cat# D8906), 1 mg/ml Escherichia coli tRNA (Sigma-Aldrich, cat# R8759), 2 mM ribonucleoside vanadyl complex (NEB, cat# S1402), and 0.02% RNase-free BSA (Invitrogen, cat# AM2618) in 2X SSC. For Stellaris MALAT1-Quasar670 probes (Stellaris, cat# SMF-2046-1), 1.25μM final concentration was used instead of the recommended 0.125μM. Probe solution in hybridization buffer were applied to coverslips followed by incubation at 37°C for 15 hours in a humid chamber. Two 30 min washes with 10% formamide in 2X SSC at 37°C were followed by post-fixation with 3.6 % paraformaldehyde in CMF-PBS buffer for 10 min at RT, followed by 3 x 5 min washes with CMF-PBS.

Coverslips were incubated with blocking buffer containing 1% BSA (RNase-free, Invitrogen, cat# AM2618), 0.5% Triton X-100 and 20μM ribonucleoside vanadyl complex (NEB, cat# S1402) in CMF-PBS for 30 min at RT. Coverslips were then rinsed three times with CMF-PBS, followed by incubation with the appropriate antibody titer (Supplementary Table 1) diluted in CMF-PBS with 0.1% Triton X-10 containing 0.4 U/μl Ribolock RNase inhibitor (Thermo Fisher Scientific, cat# EO0381) for 45 min-1 hour at RT in a humid chamber. Coverslips were then washed 3 x 5mins with 0.1% Triton X-100 in CMF-PBS for 5 minutes each. Secondary antibody incubation was performed similar to the primary antibody incubation except that the fluorescently tagged Goat anti-Rabbit / Goat anti-Mouse secondary antibodies (Supplementary Table 1) were diluted at 1:500. Subsequently, coverslips were washed 3 x 5 mins in CMF-PBS with 0.1% Triton X-100 and mounted using Mowiol-DABCO antifade mounting medium.

### Wide-field fixed-cell imaging

Wide-field images were acquired (0.08 um pixel size) using an OMX V4 microscope (Applied Precision) equipped with a U Plan S-Apo 100×/1.40-NA oil-immersion objective (Olympus), two Evolve electron-multiplying Charge-Coupled Device cameras (Photometrics). Images were aligned and deconvolved using the conservative deconvolution option within the Softworx program (Applied Precision).

### STORM imaging

For 2-color STORM imaging, we used the U2OS cell line expressing mEOS3.2-DDX39B from the endogenous locus as described elsewhere (Kim et al., 2025). Cells were fixed with 3.6% paraformaldehyde in CMF-PBS for 15 mins at RT, washed 3 x 5 mins in CMF-PBS, and then incubated in blocking buffer (3% BSA (Thermo Scientific, cat# 37520), 0.5%Triton X-100 in CMF-PBS) for 30 mins at RT, followed by 3 x 5 min washes in CMF-PBS. Coverslips were incubated overnight at 4°C with the anti-THRAP3 primary antibody (Santa Cruz Biotechnology Inc, cat# sc-133250) at 1:300 dilution in 1% BSA in CMF-PBS.

Coverslips were then washed 3 x 5 mins in CMF-PBS and then incubated for 2 hours at RT with the goat anti-mouse Fab (Jackson ImmunoResearch, cat# 115-007-003) labeled with Alexa 647 at 5μg/ml at final concentration in 1% BSA in CMF-PBS.

The Fab fragment was labeled with Alexa 647 (Invitrogen, Cat A20006) as described elsewhere (Hermanson, 2013; Bates et al., 2013). After this secondary antibody staining, coverslips were washed 3 x 5 mins in 0.1%Triton X-100 in CMF-PBS and then washed 2 x 5mins in CMF-PBS.

The STORM imaging microscope system was described elsewhere (Kim et al., 2025). Alexa 647 signals were collected first, under continuous excitation with 647 nm laser (2kW/cm^2^) and brief manual illumination with 405 nm LED illumination (0.1-0.5% intensity, Aura Light, Engine) for dye reactivation. Imaging was performed using a previously described imaging buffer (Bates et al., 2007), containing 5% glucose (Sigma-Aldrich, cat# G8270), 1% Glox (0.5mg/ml glucose oxidase, Sigma-Aldrich, cat# G2133), 40mg/ml catalase (Sigma-Aldrich cat# C100, in 50mM Tris with 10mM NaCl and 100 mM beta-mercaptoethanol (BME, Sigma-Aldrich, cat# M6250)). mEOS3.2 signals were then acquired under continuous excitation with 561 nm laser (1.5kW/cm^2^), and with brief manual illumination with 405 nm LED illumination for dye reactivation.

Images were acquired at 15-20ms per frame in continuous mode, for a total of 30000∼50000 frames. Localization, drift correction, and chromatic alignment were performed using SMAP (Li et al., 2018; Ries, 2020).

### Live cell imaging

#### Rapid wide-field imaging for short-term dynamics (second-timescale)

Cells were plated (∼48 hours before imaging) on uncoated glass bottom 35 mm dishes with 14mm diameter and 1.5 thickness coverslip (MatTek, cat# P35G-1.5-14-C) such that cells were ∼80-90% confluent at the time of imaging. For cell lines expressing proteins tagged with HaloTag, conditioned media containing Janelia Fluor(R) 646 HaloTag(R) Ligand (Promega, cat# GA1120) diluted 1:5000 was added for 1-2 hours before imaging.

For live-cell imaging we used the OMX V4 microscope together with a live cell incubator chamber (Applied Precision) and system including separate temperature control for the objective lens and incubator chamber and a humidified, 5% CO2 air supply. Both the objective lens and incubator heaters were set to 37°C; after these temperatures were reached, the cell culture dish was placed on the microscope and cells were allowed to acclimatize to these conditions for ∼1hr before beginning imaging.

Imaging was performed using simultaneous solid-state illumination with 31.3% transmittance for THRAP3/DDX39B and 10% transmittance for SON using 10-15 ms exposure for each Z-section. Optical section z-stacks (Z spacing 200 nm, ∼ 1 micron z-depth) were acquired every 1 sec for 1-2 mins.

#### Wide-field imaging for long-term dynamics (multiple hour-timescale)

Live cell imaging for greater than 10 hours was performed as described elsewhere (Kim et al., 2025) using a custom-built microscope setup.

#### 5-Ph-IAA induced SON-SRRM2 degradation

SON and SRRM2 double knockdown in the HCT116 mAID-Clover-SON SRRM2-mAID-mCherry cell line was achieved by adding to the media 5-phenyl-1H-indole-3-acetic acid (5-Ph-IAA, Tocris Biosciences, cat # 7392) to a final concentration of 5μM using a stock solution of 10mM in dimethyl sulphoxide (DMSO, Avantor cat# IC0219605590).

For live cell visualization of DDX39B during SON and SRRM2 DKD, the HCT116 mAID-Clover-SON SRRM2-mAID-mCherry cell line was transfected with the Halotag-DDX39B plasmid (Supplementary Table 1) and selected in cell culture media containing 450 μg/ml G418 (Gibco,cat# 10131035) for ∼ 2 weeks. Cells expressing Halotag-DDX39B were stained overnight with JF646 (50 pM Promega cat# GA1120). After adding 10∼20μM 5-Ph-IAA, we acquired live cell movies to observe SON and SRRM2 degradation while visualizing Halotag-DDX39B.

#### RNP transgene induction and RNP localization relative to DDX39B, THRAP3, and SON / SRRM2 (NS)

U2OS cell lines stably expressing the E6 β-globin and mini-Dystrophin A-1 transgenes tagged with the MS2 repeats and expressing a YFP-fused MS2 binding protein (Mor et al., 2010; Brody et al., 2011), were seeded (∼30,000 cells) on 12mm #1.5 coverslips (Electron Microscopy Sciences cat# 72230-01). For E6, cells were treated with 15 µg/ml doxycycline (Sigma-Aldrich cat#D5207) for 6, or 16 hours, while mini-dystrophin cells were treated with 1 µg/ml Ponasterone A (Enzo Life Sciences) for the same durations to induce transgene expression. Cells were fixed in 4% PFA for 20 mins at RT. Cells were permeabilized in 0.5% Triton X-100 in 1x PBS for 10 min, and blocking was applied using a buffer composed of 0.1% Triton X-100 and 5% bovine serum albumin (BSA) fraction V (MP Biomedicals cat# 160069) for 1 hour at RT. Primary antibodies were diluted in blocking buffer (Supplementary Table 1) and applied to coverslips overnight at 4°C for 16 hours in a humidified chamber. Coverslips then were washed for 5 mins three times using 0.1 % Triton X-100. This was followed by incubation with secondary antibodies for 1 hour at RT and then washes for 5 mins three times using 0.1 % Triton X-100. The nucleus was stained with SYTOX deep red nucleic acid stain (Invitrogen cat# S11381) for 20 min, followed by final rinses with 1x PBS. Secondary antibodies used; Alexa Fluor 405-conjuated goat anti rabbit IgG (1:500, Abcam cat# ab175652) and Rhodamine red-conjugated donkey anti mouse IgG (1:250, Jackson Immunoresearch cat# 715-297-003).

Confocal imaging was performed on a Leica SP8 inverted microscope. The objective used was 100x 1.4 NA, with Leica immersion oil, at room temperature. The mounting medium was custom made 80% glycerol with p-phenylenediamine antifade (Sigma). The resulting images were deconvolved with Huygens Professional (Scientific Volume Imaging, Hilversum, The Netherlands) using the CMLE algorithm, with signal-to-noise ratios of 12 and 5 iterations, using the Huygens confocal module.

### RNA-Seq sample preparation and analysis

#### Cell synchronization and protein depletion

For synchronization at the G1/S boundary (JavanMoghadam-Kamrani and Keyomarsi, 2008), HCT116 mAID-Clover-SON SRRM2-mAID-mCherry CMV-OsTIR1(F74G) cells were seeded in 6-well plates and grown to approximately 80% confluency, followed by treatment with 20 µM Lovastatin (Mevinolin; LKT Laboratories, cat# M1687) for 24 hours. To release the G1 arrest but block cells at the G1/S boundary, cells were rinsed three times with pre-warmed, CO₂-equilibrated culture medium supplemented with 2 mM Mevalonic acid (Mevalonolactone; Millipore-Sigma cat# M4667) and 2 mM thymidine (Millipore-Sigma, cat# T9250).

Degradation of AID-tagged proteins was induced by the addition of 5 µM 5-Ph-IAA simultaneously with the release of the G1 block by addition of the Mevalonic acid/thymidine-containing media. Cells were incubated for 16 hours after this G1 block release. Control cells were treated the same except that an equivalent volume of DMSO without added 5-Ph-IAA was added. The G1 arrest followed by addition of 5-Ph-IAA coincident with the release of this block for the G1/early S synchronization was used to minimize cell cycle-related variability between DMSO-treated controls and the SON/SRRM2-depleted samples.

#### RNA extraction and RNA-seq

Cells in a 6-well plate were rinsed twice with ∼1 mL of cold CMF-PBS per well, and total RNA was extracted immediately using the RNeasy Plus Mini Kit (Qiagen, cat# 74134), following the manufacturer’s protocol.

Purified RNA eluted in RNase-free water was treated with 2U DNase I (New England Biolabs, cat# M0303S) for 20 mins at 37°C, followed by DNase I inactivation by adding 5 µM EDTA (Thermo Fisher Scientific, cat# J15694.AE) and 10 mins incubation at 75°C.

RNA integrity was assessed at the DNA Services Facility, Roy J. Carver Biotechnology Center (University of Illinois at Urbana-Champaign), using an Agilent Bioanalyzer. All samples exhibited RNA Integrity Numbers (RIN) of 10. Ribosomal RNA was depleted using the FastSelect HMR kit (Qiagen, cat# 334386). RNA-seq libraries were prepared with the KAPA HyperPrep Stranded mRNA Library Kit (Roche, cat# KK8580), quantified by qPCR, pooled, and sequenced on a NovaSeq X Plus instrument using V1.0 sequencing chemistry for 151 paired-end cycles. Adapter trimming was performed on the 3′ ends of reads, using the first 33 nucleotides of the adapter sequences for trimming. FASTQ files were generated and demultiplexed using Illumina’s bcl2fastq v2.20 software.

#### RNA-seq analysis

The quality of trimmed reads was assessed using FastQC with default parameters. High-quality reads were aligned to the human reference genome (GRCh38/hg38) using STAR aligner (Dobin et al., 2013), with the corresponding GTF annotation file obtained from Ensembl. Differential gene expression analysis was performed using the DESeq2 R package (Love et al., 2014) using DMSO-treated samples as controls. Genes with an adjusted p-value (Padj) < 0.05 (Benjamini-Hochberg correction) were considered significantly differentially expressed (Supplementary Table 2).

### Image analysis

All fixed-cell images shown are single optical sections unless indicated otherwise. Non-linear exponential gamma scaling of 0.8 was applied to all images of DDX39B, MALAT1 and RBM8A to reduce the intensity dynamic range and thus facilitate the visualization of low-intensity nuclear regions. Images presented in the live-cell imaging videos are 2D maximum-intensity projections of the original optical section z-stacks acquired at each time point. All images were processed using Fiji (ImageJ) (Schindelin et al., 2012).

### Protein-intensity quantification within and outside NS

A single optical z-section containing the max-signal over the entire z-stack for NS (SON or SRRM2) was selected using the np.max function in the Python NumPy package (Harris et al., 2020). The intensity distribution of each candidate protein inside versus outside of NS was assessed as follows: Pixels with intensities < 10% of the maximum signal of the protein were considered “noise” and discarded. A nuclear mask, established by DAPI staining, was applied to both the NS and candidate protein images. NS were defined by SON/SRRM2 intensity thresholds established using Otsu thresholding (Otsu, 1979). The remaining pixels showing candidate protein “positive” signals (>10% maximum intensity threshold) were then divided into two categories: “Protein within NS” for pixels overlapping with the NS segmented regions and “Protein outside NS” for remaining positive pixels outside the NS segmented regions. The resulting images were displayed using the “viridis” LUT color-map. Cumulative-intensity percentile plots of the “Protein within NS” and “Protein outside NS” staining distributions were generated using the Python package Matplotlib (Hunter, 2007).

### Network changes upon transcriptional inhibition with DRB

A Fiji macro (“Skeletonize”, See Code Availability Section) was created for this quantification. Briefly, for each image, the SON channel was used to generate a NS mask. Then, all pixels of the protein channel that overlapped with the NS mask were set to zero intensity to generate an image of only the protein outside NS. The resultant image was converted into a binary image using Otsu thresholding (Otsu, 1979). Using the Fiji Skeletonize function (fiji.sc/Skeletonize3D), “protein skeletons” were generated, and branch lengths and number of branches/skeleton were calculated. Total skeleton length was computed by summing the lengths of all the skeletons within a nucleus. Number of connections was the number of unique skeletons (length > 5 pixels) identified within a single nucleus.

### Distance measurement for RNA Pol II foci relative to SON, THRAP3 and DDX39B networks

RNA Pol II Ser2p and Ser5p staining foci locations were determined using the ‘Find Maxima’ function in ImageJ. Detected locations were assigned a pixel value of 1, while all other pixel values were set to 0, converting the Pol II image into a binary image containing only Pol II locations. The binary image was then imported into a custom MATLAB code for distance measurements relative to NS (https://github.com/Jiahkim/distancebtwlocusNS.git) . Pol II-positive pixels were first classified based on whether they overlapped with NS regions. NS areas were defined by segmentation of a target NS marker using local thresholding (Bradley and Roth, 2007). For Pol II-positive pixels outside NS area, distances to the nearest pixel at the NS boundary were as calculated.

### Distance measurement for Type I and Type II DNA-FISH spots relative to SON, THRAP3 and DDX39B

To measure distance of a spot to SON, THRAP3 and DDX39B, we used the Loci_Compartment_Distance ImageJ plugin described elsewhere (Kumar et al., 2024). Since the THRAP3 and DDX39B networks are widely distributed within the nucleus, and due to limited Z-resolution, we used a single optical section for THRAP3 and DDX39B analysis. For NS detection, we used the sum-projection of 3 optical sections centered over the brightest spot of the locus. To define the edge of the compartment boundary, the intensity drop of 50% of the local maximum intensity (after background correction) was used to locally segment the image. The center of mass of the DNA FISH spot signal was used as the spot center location and the shortest distance between the spot center to the nearest compartment edge was measured.

### Categorizing and assessing alignment of DNA-FISH trajectories relative to SON, THRAP3 and DDX39B foci

To eliminate DNA FISH background staining signals, for the DNA FISH mapping of chromosome trajectories, only the “Apex” biotin signals that colocalized adjacent to DNA-FISH signals for the “Arm” regions were considered as true positive Apex signals. Additionally, the Apex signal had to be at least 2-fold higher intensity than single biotin spots seen elsewhere in the nucleus to be considered for analysis. For analysis, a sum-projection was generated of all the consecutive optical sections which contained an individual trajectory FISH signal consisting of positive DNA-FISH Apex and Arm signals. Chromosome trajectories were categorized as condensed if the Arm region appeared as a single spot or blob next to an Apex DNA-FISH spot. Any Arm region appearing as 2 or more spots was termed “decondensed”. For alignment, a trajectory was considered aligned only if all the spots of the Arm and Apex were either overlapping or adjacent and in-contact to THRAP3 or DDX39B foci. In all other cases, the chromosome trajectory was classified as not aligned.

### Distance measurements of Trajectory Apex versus Arm regions from nearest NS, THRAP3, or DDX39B foci

DNA-FISH spots for Trajectory Apex and Arm were detected using the “blob log” function within the scikit-Image Python package (Van Der Walt et al., 2014) and were clustered using DBSCAN (Ester et al., 1996). For this clustering, an iterative threshold estimation method was employed such that the number of clusters per nucleus was 3-4. For each cluster, the cluster center was calculated and a box of 30x30 pixels (x-y) from the center was cropped; within this cropped volume, 7 optical sections (z-slices) were sum intensity projected to create a new 2D image, within which the center of mass of the Apex signal was calculated. Within the same 2D cropped projection, the Arm region signal was segmented and the pixels within the segmented region were split into three categories, for each of which their center-of-mass determined: 1) “Arm-close”: 25% pixels closest to Apex; “Arm-far”: 25% pixels farthest from Apex; “Arm-middle”; remaining pixels. The DDX39B or THRAP3 and NS signals were thresholded using the Otsu function (Otsu, 1979). The shortest Euclidean distance between the center of each DNA-FISH spot (Apex or Arm) to the nearest edge of the segmented protein/speckle was measured.

### Distance measurements of E6 and mini-Dystrophin transgene RNAs relative to SON, THRAP3 and DDX39B foci

Distance measurements for transgene RNAs, tagged with MS2 repeats and MS2-binding proteins, relative to NS (SON/SRRM2), DDX39B, or THRAP3 foci were performed similarly to the RNA Poll II foci distance measurements. Nuclear regions were manually selected, and RNA spots within each nuclear region were identified using the ‘Find Maxima’ function, to generate binary images. NS areas were segmented using the Otsu’ thresholding algorithm (Otsu, 1979) which effectively defined perispeckle and NS boundaries. For this data, plotting and statistical tests were performed using GraphPad Prism.

### Distance measurement of HSPH1 and HSPA1 RNAs relative to SON, THRAP3, and DDX39B foci

For HSPH1 and HSPA1 RNA-FISH analysis, images were denoised using the “despeckle” function in ImageJ and a Gaussian filter of 1 pixel radius applied afterwards Distance measurements relative to SON, THRAP3 and DDX39B foci were performed using the ImageJ plugin as described for DNA-FISH measurements of Type I and Type II SON TSA-seq peaks.

For Mander’s coefficient calculations, we used a maximum-intensity projection of 3 optical sections of the “despeckle” denoised and Gaussian fitered images. A nuclear mask was generated using the DAPI signal and Otsu thresholding (Otsu, 1979) and applied to the RNA and protein signals. These nuclear-masked images were then used for calculating Mander’s coefficient using the JaCoP plugin (Bolte and Cordelières, 2006) in Fiji (Schindelin et al., 2012).

### Plotting and Statistical Analysis

All plots, unless otherwise stated, were generated using the ggplot2 package in R version R4.2.2. All analysis of variance (ANOVA) used type 3 comparison using the “car” package. The options were set as options (contrasts=c (“contr.sum”, “contr.poly”)). We used a generalized linear model with gamma distribution followed by the Tukey posthoc test. For RNA Pol II foci and RNA-FISH distance measurements, the Kruskal-Wallis test followed by the pairwise Wilcoxon test was used to calculate P-values as indicated in the figures. For Mander’s coefficient comparison, an unpaired t-test was used in R. All boxplots represent the interquartile range with the upper line indicating 75% and the bottom line indicating 25%. Central line indicates median, and whiskers represent the range of data shown.

## Supplementary Figure Legends

**Supplementary Figure 1: Screening for proteins exhibiting perispeckle patterns**

A: Representative protein candidates showing different categories of distributions within the interchromatin space (ICS). (Left to right) SRRM2: NS-enriched protein with no extended staining network in ICS; DDX39B: NS-enriched protein with additional extended staining network in ICS; protein concentration is highest within NS and lower in ICS network than within NS; THRAP3: protein concentrations are similar in and/or surrounding NS as the extended staining network in ICS; EIF4A3: protein concentration is present within NS but protein has a diffuse distribution, rather than a network-like staining pattern, within ICS; SAF-B: protein is largely absent within NS but shows an extended staining network not obviously correlated with NS within ICS. Each protein distribution is shown in grey (top row) and relative to NS (SON or SRRM2, magenta) (top 2 rows; scalebar: 5 μm). Regions highlighted within the whole nuclear images magnified to show protein distribution (grey) relative to NS (magenta; 3rd and 4th rows; scalebar: 1 μm). Pseudo-color images show the protein distribution segmented into “protein within NS” and “protein outside NS” fractions. Intensity plots (bottom row) show protein intensities over every percentile for “protein within NS” (blue) and “protein outside NS” (orange) fractions.

B: Plot showing fraction of total protein intensity found outside NS. (n (number of nuclei) =3). Each point represents mean± standard deviation (SD).

C: Plot showing fraction of nuclear area occupied by protein distribution outside NS (n (number of nuclei)=3). Each point represents mean±SD.

D: Representative images of protein distributions all candidates surveyed in the screening ordered based on fraction of protein staining outside of NS (low to high). Pseudocolor intensity-mapped images (middle and right panels) highlight the lower intensity protein distributions seen within the ICS.

E: Protein distributions segmented into “Protein within NS” and “Protein outside NS” intensity-mapped images for each candidate.

F: PLRG1 colocalizes with THRAP3 in staining network outside of NS but is enriched within NS: Immunostaining of PLRG1(gray) along with endogenously tagged mCh-THRAP3 (magenta) U2OS cells. (Main image-scalebar: 5 μm; inset-scalebar: 1 μm). G: Cells expressing lentiviral mediated mCherry MAGOH (gray) immunostained with DDX39B (green) show colocalization of both proteins within ICS. (Main image-scalebar: 5 μm; inset-scalebar: 1 μm).

H: RNA FISH (using oligonucleotide probes) distribution for MALAT1 (grey) relative to THRAP3 (magenta) and DDX39B (green) immunostaining within the ICS. Second and fourth rows represent enlargements of boxed regions (white borders) from first and third rows, respectively. Apart from NS-enriched MALAT1, diffraction-limited spots of MALAT-1 are seen in the ICS in the vicinity of NS and show close spatial relationship to THRAP3 and DDX39B foci. Box plot showing fraction of RNA signal outside NS overlapping with THRAP3 or DDX39B signals. (n (number of nuclei) =10 for each comparison). White square represents mean and central line represents median value.

**Supplementary Figure 3: Transcriptional inhibition causes changes in perispeckle network PSTP.**

A-B: Disruption of THRAP3 (magenta) network upon transcriptional inhibition also seen with Triptolide (A) and α-amanitin (B) treatments. NS marked by SON in yellow. The extent of disruption of the THRAP3 pattern after 2 hours of Triptolide and α-amanitin treatment resembles the disruption observed with 1 hour DRB treatment.

C: Intensity-mapped images of THRAP3 (1st column) and DDX39B (3rd column) and their corresponding skeletons (2nd and 4th columns) upon treatment with 5,6-Dichloro-1-β-D-ribofuranosylbenzimidazole (DRB) comparing changes with DMSO control versus 1 and 2 hours (h) of treatment.

D: Multiple members of PSTP colocalize even as the networks are disrupted. Comparison of BCLAF1 (immunostaining, grey) and EGFP-THRAP3 (endogenous gene tagged, magenta) networks shows colocalization of BCLAF1 and THRAP3 as they are both disrupted upon DRB treatment (2 hours (h)). NS are marked by HaloTag-SON (endogenous gene tagged, yellow). Scale bar-5 μm main image; 1 μm inset.

E-F: Intensity mapped images and their corresponding skeletons of BCLAF1 (PSTP member) and RBM8A (PSDP member) upon DRB treatment as compared to DMSO shows similar disruption of PSTP network versus modest perturbation of PSDP network using staining against other PSTP (BCLAF1) and PSDP (RBM8A) components.

G-H. Boxplots showing total skeleton length (G) (in pixels) and number of connections

(H) of BCLAF1 (left, pink and magenta) and RBM8A (right, light and dark green) with DMSO or DRB treatment for 2 hours. n (number of nuclei) = 10-13 for each condition. One-way ANOVA with Tukey post-hoc test was used to calculate P-values. ns : P > 0.05; * : P ≤ 0.05; ** : P ≤ 0.01; *** : P ≤ 0.001; **** : P ≤ 0.0001.

**Supplementary Figure 4: RNA Polymerase II foci associate closely with PSTP as compared to PSDP**

A-B: Representative images of RNA Pol II foci (Ser5p top rows, Ser2-bottom rows) detected for distance measurements from THRAP3 (magenta, A) and DDX39B (magenta, B) and SON (green, B). Shown alongside are line profile plots of RNA Pol II foci relative to the three compartments. RNA Pol II foci appear to frequently localize adjacent to or overlapping with staining concentrations of THRAP3 and DDX39B, as seen by offset of their peaks with RNA pol II foci. Images on the right show segmented THRAP3 (top) and DDX39B (bottom) networks used for distance measurements of RNA Pol II foci. Circled foci colored based on distance: Red: <0.15 μm; Blue: >0.15 μm and <0.45 μm; Yellow: ≥0.45μm.

**Supplementary Figure 5: TSA-seq reveals gene expression hot-zones preferentially associate with PSTP and PSDP also in U2OS cells**

A: Representative TSA-seq and Lamin B1 DamID tracks for chromosome 2 in U2OS cells: SON (blue), DDX39B (green), THRAP3 (magenta) and Lamin B1 DamID (grey). Yellow highlights mark Type I peaks and blue highlights mark Type II peaks.

B: Heat map showing Pearson’s correlation coefficients for SON, THRAP3 and DDX39B TSA-seq datasets (mean of two replicates) in U2OS cells.

C: DNA-FISH (yellow) for Type I and Type II peaks (Apex) relative to SON (cyan) and THRAP3 (magenta) or DDX39B (green) and SRRM2 (blue). Type I regions are NS-associated and remain in contact with THRAP3 and DDX39B. Type II regions that are away from a NS are also in contact or close proximity of THRAP3 or DDX39B accumulations. Regions highlighted in white boxes are shown on the right as insets. Scale bars-5 μm main image; 1 μm inset.

D: Violin plots showing distance of DNA-FISH spots relative to SON, DDX39B and THRAP3 in U2OS cells. Central line indicates median distance and white square within the boxplot indicates mean distance. (n (alleles) as indicated: 126 (Type I) and 104 (Type II) SON; 130 (Type I) and 106 (Type II) DDX39B; 126 (Type I) and 104 (Type II) THRAP3). One-way ANOVA with Tukey post-hoc test was used to calculate P-values. ns : P > 0.05; *** : P ≤ 0.001; **** : P ≤ 0.0001.

E-G: Probability density plots for distance distribution of DNA-FISH spots relative to SON (E), DDX39B (F) and THRAP3 (G) for Type I (yellow) and Type II (blue) peaks in U2OS cells. (n (alleles) as indicated: 126 (Type I) and 104 (Type II) SON; 130 (Type I) and 106 (Type II) DDX39B; 126 (Type I) and 104 (Type II) THRAP3).

**Supplementary Figure 8: NS knockdown does not alter spatial positioning of genome relative to DDX39B and THRAP3 in HCT116 cells**

A: Representative images showing mClover-SON (green) and SRRM2-mCherry (magenta) fluorescence intensities change over time with 5-Ph-IAA treatment for 0 min, 2 hours or 4 hours.

B: Heat map showing Pearson’s correlation coefficients across WT HCT116 (SON, THRAP3, DDX39B) and HCT116 SON-SRRM2 degron cells (DMSO THRAP3, 5-Ph-IAA THRAP3, DMSO DDX39B, 5-Ph-IAA DDX39B) TSA-seq datasets.

C: TSA-seq score tracks for chromosome 2 in HCT116 SON-SRRM2 degron cells: SON (TSA-seq score from WT cells; black shown for comparison) Type I (yellow highlights) and Type II Peaks (blue highlights). Normalized TSA-seq scores between DMSO-treated and 5-Ph-IAA-treated cells show small changes all over the entire chromosome for THRAP3 (magenta; negative residual; P-value in grey significant) and DDX39B (green; positive residual; P-value in grey not significant). P-value threshold set at log10 scale at 2 (red line).

D: Ideogram plots of all chromosomes in percentile change (DMSO-treated minus 5-Ph-IAA treated) in TSA-seq scores in HCT116 SON-SRRM2 degron cells for DDX39B (left) and THRAP3 (right) show little or no changes (-5 to +5 green) except for very few bins showing higher changes (indigo -20 / yellow +20).

E: Scatterplots between mean TSA-seq scores of DMSO-treated and 5-Ph-IAA-treated DDX39B (green) and THRAP3 (magenta) show similar correlation coefficient as two individual replicates of DMSO-treated DDX39B (blue) and THRAP3 (yellow) TSA-seq

## Supplementary Information

Supplementary Table 1: Reagent information (Primers, antibodies, oligolibraries and related information).

1. cDNA constructs generated for screening: Primer sequences used for PCR amplification and restriction enzymes used for cloning
2. Antibodies: Details and dilutions used
3. CRISPR-Cas9 mediated gene tagging: Target specific gRNA sequences, homology arm sequences for Donor template generation and genotyping primers used.
4. Selected Type I and Type II genomic regions for FISH validation of TSA-seq in HCT116 and U2OS cells
5. Oligolibrary DNA-FISH probe sequences for 7 Type I peaks
6. Oligolibrary DNA-FISH probe sequences for 8 Type II peaks
7. Genomic coordinates of selected regions of Trajectory FISH of TypeI peak to nearest LAD or Type II peak to nearest LAD region
8. Oligolibrary DNA-FISH probe sequences for Apex of Type I peak
9. Oligolibrary DNA-FISH probe sequences for Arm regions of Type I trajectory
10. Oligolibrary DNA-FISH probe sequences for Apex of Type II peak
11. Oligolibrary DNA-FISH probe sequences for Arm regions of Type II trajectory
12. Human MALAT1 oligolibrary RNA-FISH probe sequences
13. Human MALAT1 Stellaris RNA-FISH probe sequences
14. 4DN Data Portal Accession numbers of TSA-seq datasets generated for this study.
15. 4DN Data Portal Accession numbers of RNA-seq generated for this study.

Supplementary Table 2: Complete list of significantly affected genes after nuclear speckle knockdown

## Supplementary Videos

**Video 1: Rapid dynamics of THRAP3 network in U2OS**

A video of EGFP-THRAP3 (left, green) and its corresponding intensity mapping (right) in live U2OS cells showing THRAP3 non-diffuse accumulations forming, disappearing and reforming in the same space within tens of seconds. Images are acquired at 1 second intervals. Video is displayed at 5 frames per second. NS are marked by HaloTag-SON (left, magenta).

**Video 2: Dynamics of DDX39B network in U2OS cells**

A representative video of mCherry-DDX39B (left, green) and its corresponding intensity mapping (right) in live U2OS cells showing connections of DDX39B forming and disappearing in the vicinity of NS marked by EGFP-SON (left, magenta). Images are acquired at 1 second intervals. (Gamma 0.8 used to display DDX39B). Video is displayed at 5 frames per second.

**Video 3: Perispeckle THRAP3 network remains stable over many hours within the interchromatin space**

A video of mCherry-THRAP3 (magenta), EGFP-SON (yellow) and SiR-Hoechst DNA (cyan) imaged over 16 hours (h) at 40-minute intervals. THRAP3 network is dynamic but remains overall stable over time within the interchromatin space.

**Video 4: Perispeckle DDX39B network within the interchromatin space remains stable over many hours**

A video of mCherry-DDX39B (magenta), EGFP-SON (yellow) and SiR-Hoechst DNA (cyan) imaged over 13 hours (h) at 10-minute intervals. DDX39B connections fluctuate but reappear within the same space over time making the overall network highly stable.

**Video 5: Transcriptional inhibition by 5,6-Dichloro-1-β-D ribofuranosylbenzimidazole (DRB) causes gradual disruption of THRAP3 network**

A video of EGFP-THRAP3 (left, green) and HaloTag-SON (left, red) along with intensity mapping of THRAP3 (right) shows the network of THRAP3 collapses into nuclear speckles over the course of DRB treatment. Imaging was started 10 minutes (min) after adding DRB and images were taken every 2 minutes for up to 2 hours.

**Video 6: Live imaging of DDX39B upon nuclear speckle knockdown**

Live-cell imaging of transiently expressed HaloTag-DDX39B (green, center) along with intensity mapping (right) and mClover-SON (in magenta, left) and SRRM2-mCherry in HCT116 cells shows DDX39B network is maintained even after double knockdown of SON and SRRM2 and complete loss of nuclear speckles. DDX39B also shows minimal changes to “speckle-like” structures corresponding to the concentration of DDX39B in regions originally occupied by the NS identified by SON and SRRM2 concentrations.

Images were taken every 5 minutes (min) beginning 20 minutes after start of 5-Ph-IAA treatment.

## Data Availability

All the raw image data are available upon request. All TSA-seq and RNA-Seq datasets (raw and processed files) generated in this study are available at the 4DN Data Portal (see Supplementary Table 1 for ID numbers) and the GEO (GSE304818)

## Code Availability

All custom scripts generated for this study are available on Github (https://github.com/nehacv93/PerispecklePatterns_partitionICS).

## Author Contributions

N.C.V. and A.S.B. conceived of and designed the study; N.C.V. performed most experiments and analyzed data including the screening, fixed-cell imaging, TSA-seq, oligo library design and DNA-FISH. N.C.V. and S.M. performed short timescale live-cell imaging of networks and DRB treatments. J.K. performed long term live-cell imaging, STORM and image analysis for the Pol II distribution and reporter transgene RNP distances relative to the networks. G.F. performed experiments of E6 and mini-Dys transgene RNPs and STED imaging. S.M. and N.C.V. replicated the transgene data set and also performed RNA-FISH for endogenous genes. Endogenous RNA-distance measurements were by N.C.V. and S.M. performed Mander’s coefficient analysis. A.B. and M.K. generated and validated the SON-SRRM2 degron cell line. G.H., N.C.V., J.K. and S.M. made the endogenous fluorescent tagged cell lines by CRISPR-Cas9. P.C. performed FACS analysis of degron cell line and RNA-Seq analysis. N.C.V. performed TSA-seq and RNA-Seq correlation analyses. J.D. helped design and supervised the initial implementation of primary screen. K.Y.H. and Y.S.T provided reagents and supervision. N.C.V. and A.S.B. wrote the manuscript with inputs from other authors.

## Acknowledgements

This work was supported by the National Institutes of Health Common Fund 4D Nucleome Program grant U01DK127422 (ASB, KYH, YS) and also NIH R01GM058460 (ASB) and JSPS KAKENHI JP23H04925 and JP25H00979 (MTK) grants. We acknowledge assistance provided by the DNA Services, Roy J. Carver Biotechnology Center, University of Illinois at Urbana-Champaign for quality control on TSA-seq libraries, RNA-seq library construction, and sequencing (TSA-Seq and RNA-Seq libraries). Custom RNA-Seq scripts for processing and analysis of data were provided by the High-Performance Computing in Biology group (HPCBio), Roy J. Carver Biotechnology Center, University of Illinois at Urbana-Champaign. We thank Ratna Karatgi for advice regarding statistical analysis and Mishal Assif for assistance with image analysis.

