## Supplementary Figures for "Highly active chromosome regions preferentially associate with two perispeckle networks that partition the interchromatin space"

**A** **Different categories of protein distributions observed relative to nuclear speckles (NS) within the interchromatin space (ICS):**

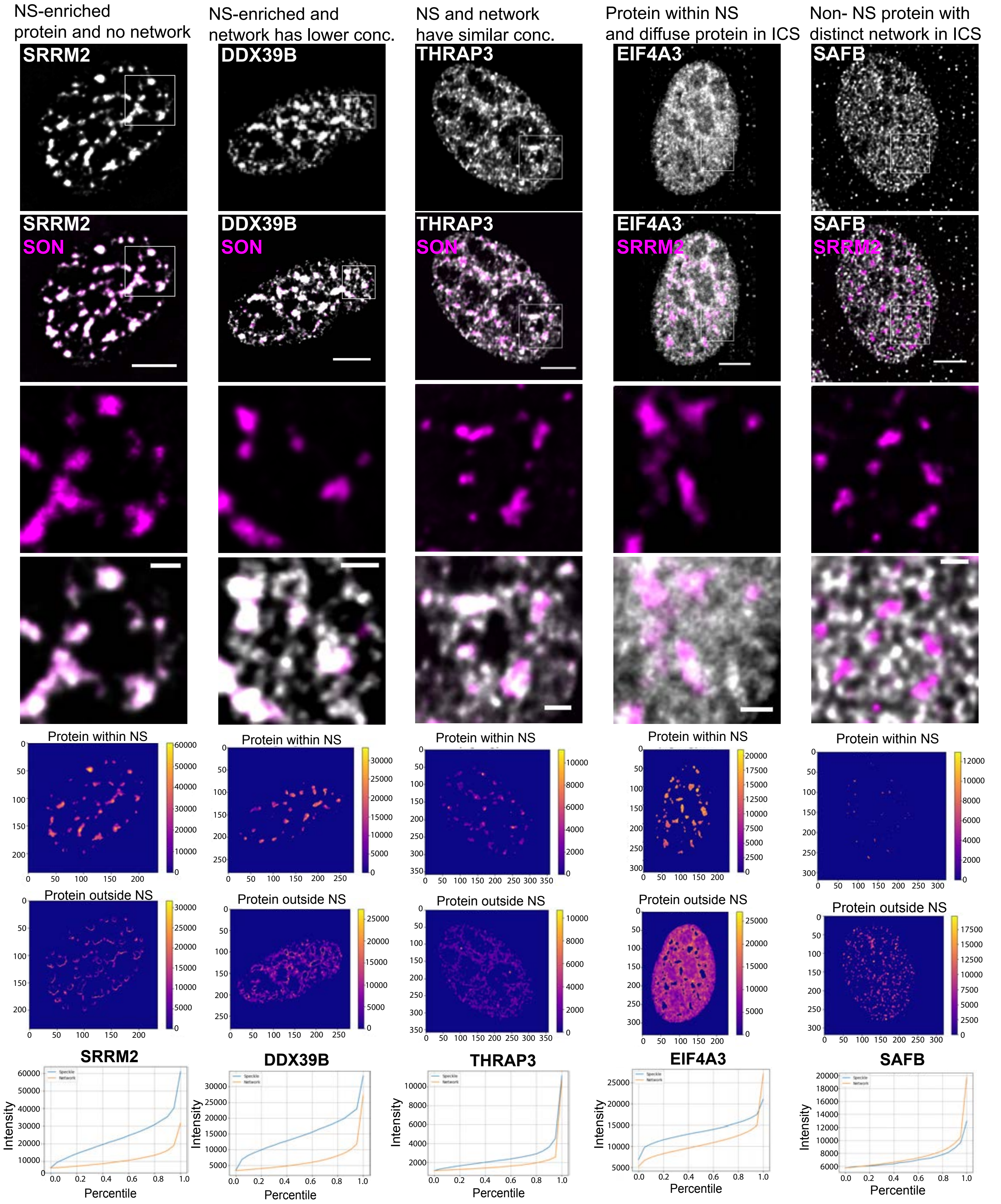


**Supplementary Figure 1**

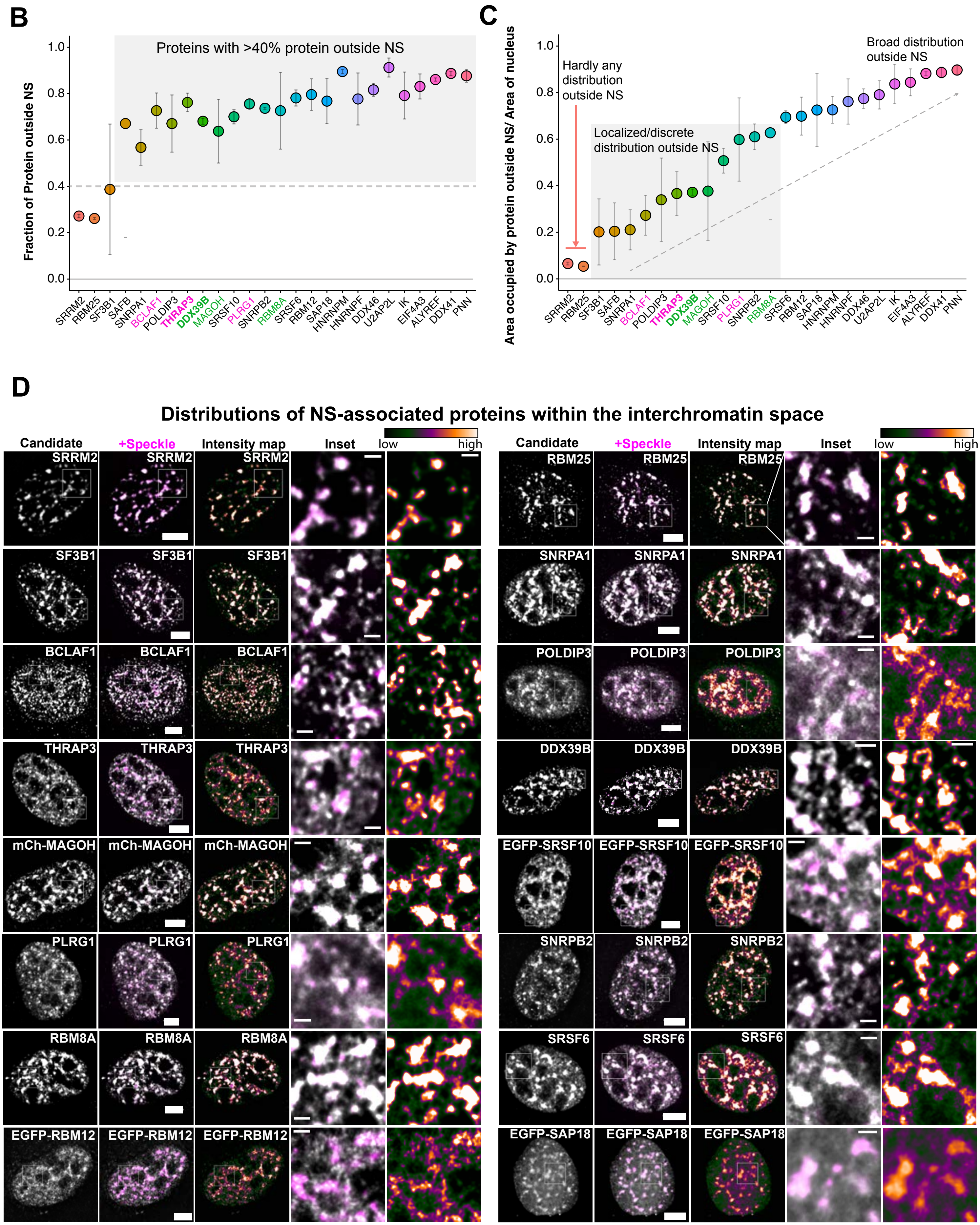


**Supplementary Figure 1**

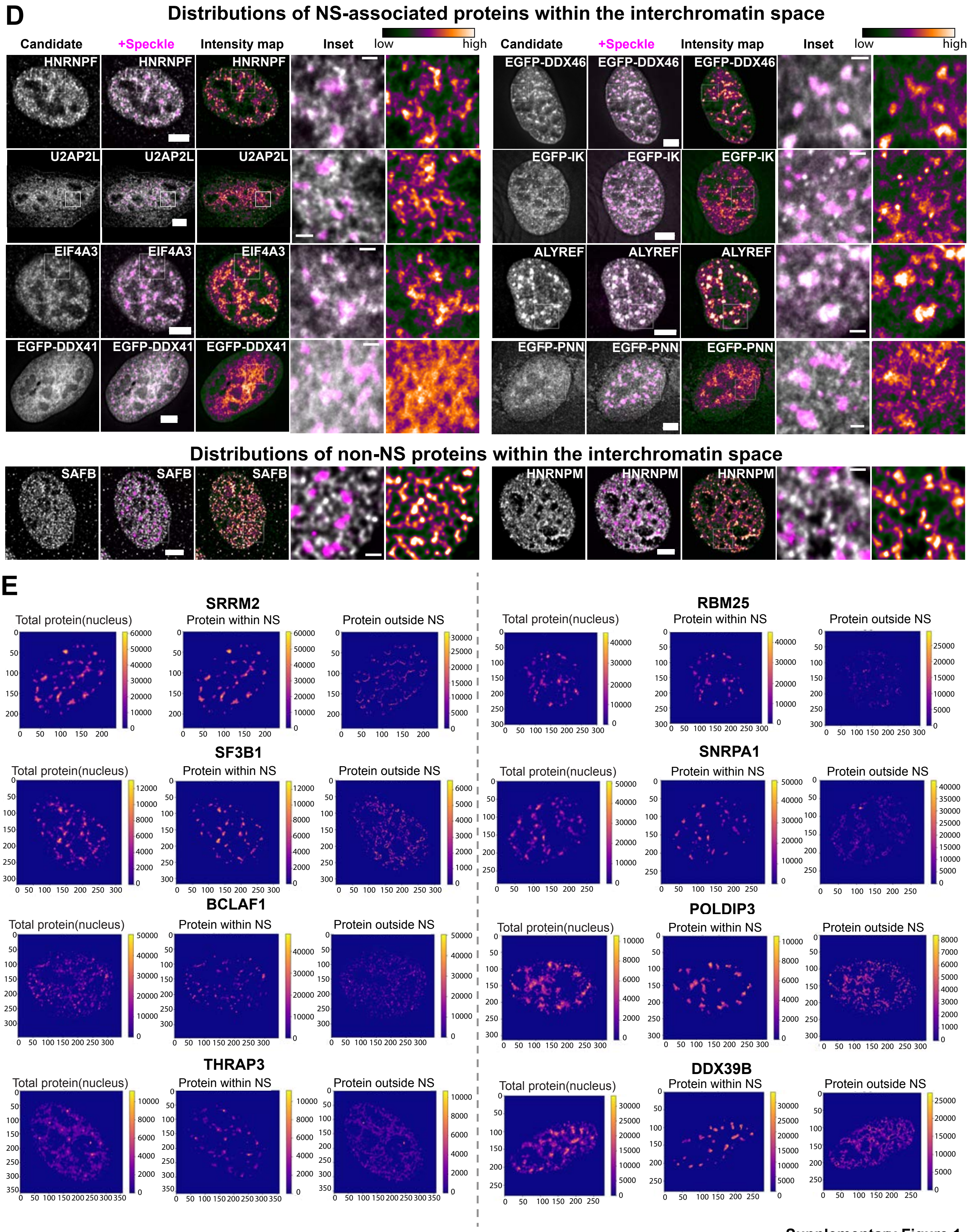


Supplementary Figure 1

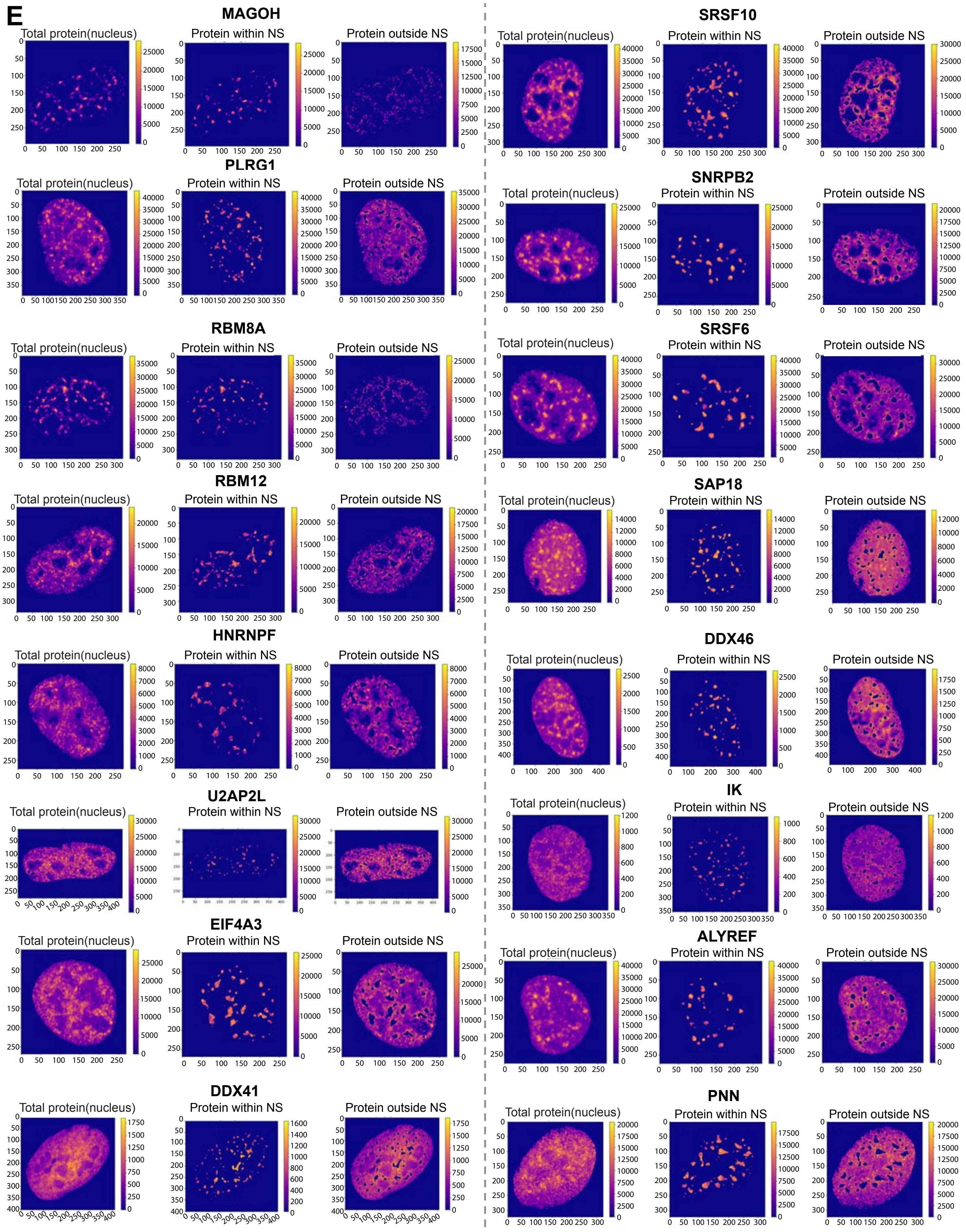


**Supplementary Figure 1**

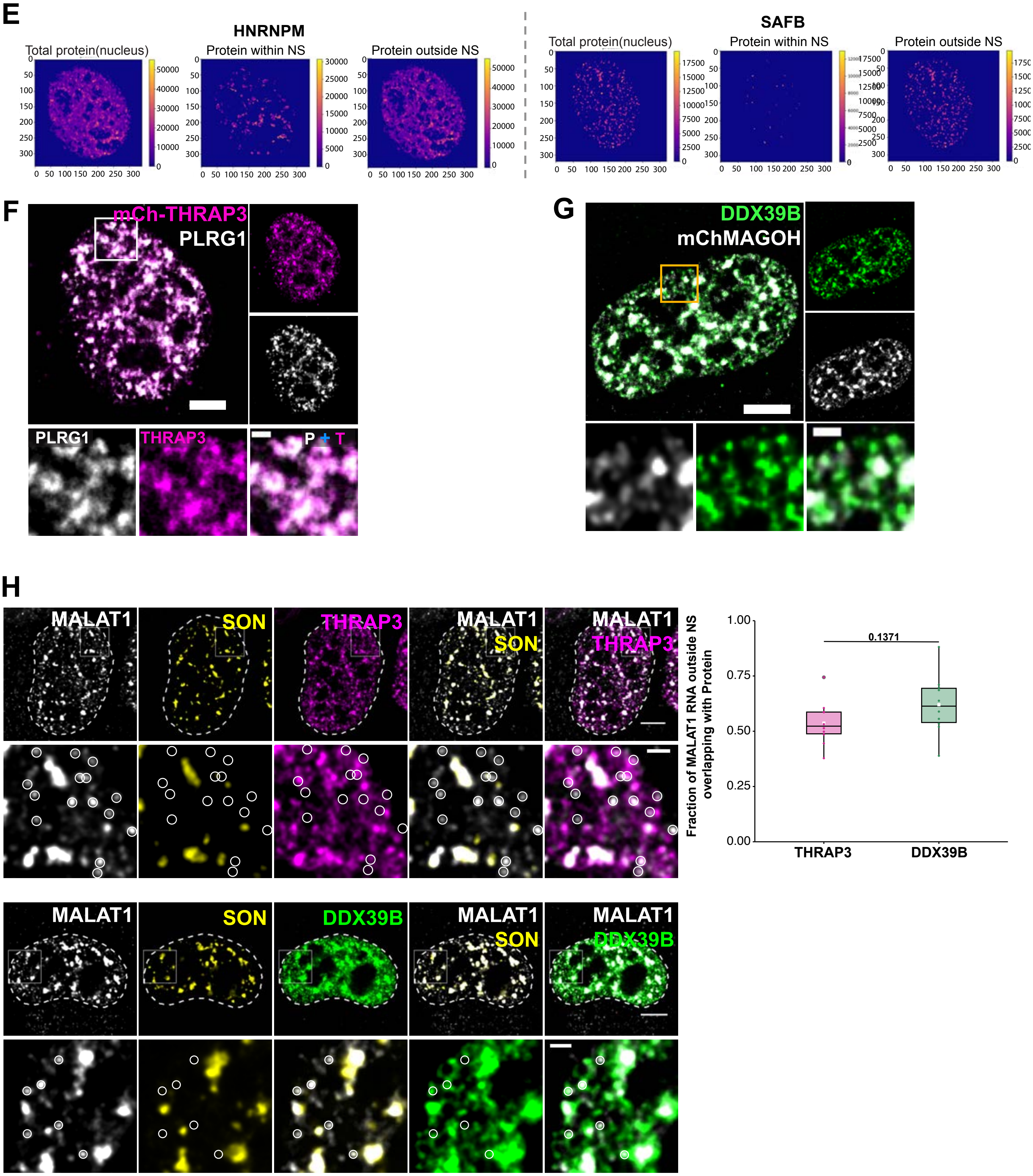


Supplementary Figure 1

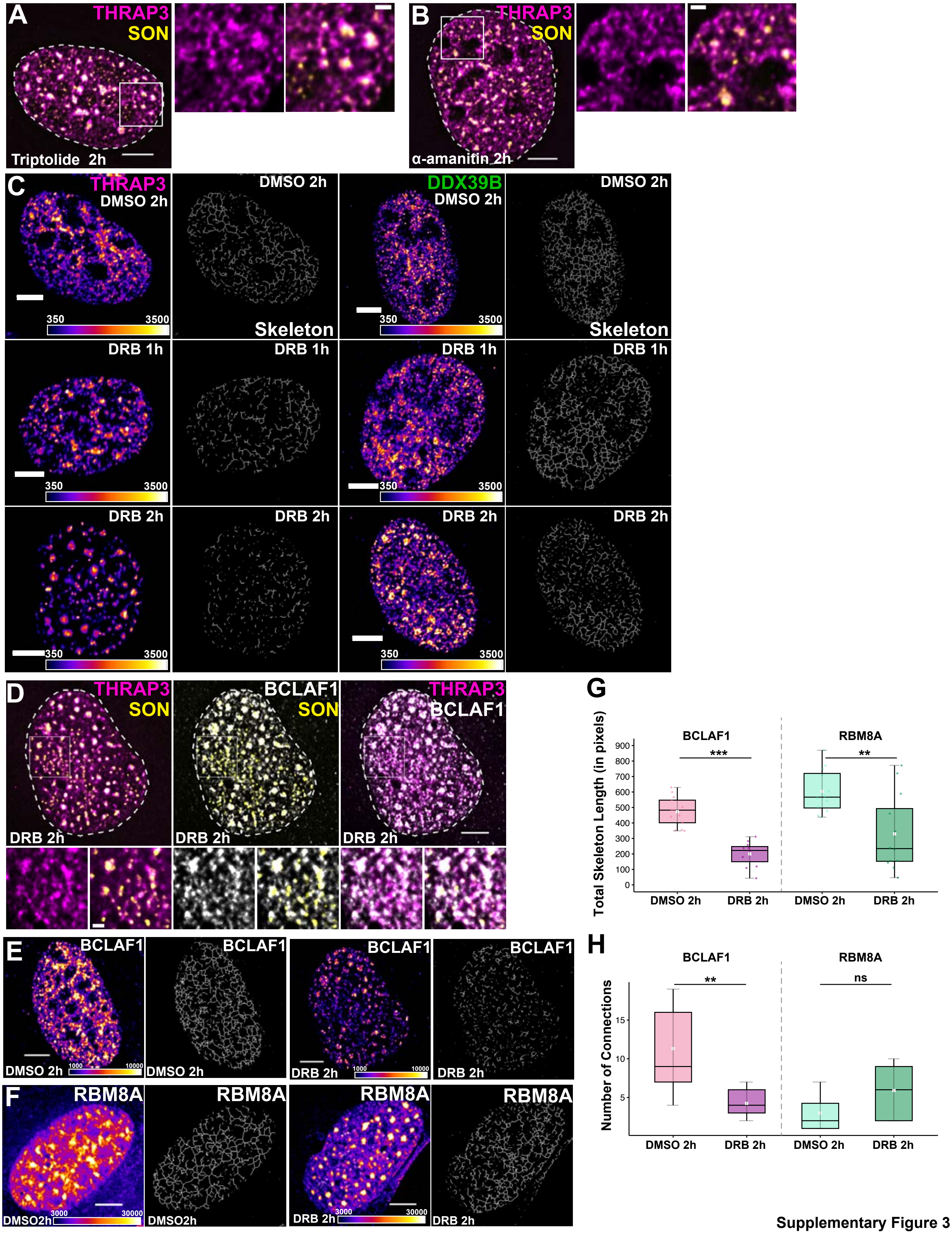


Supplementary Figure 3

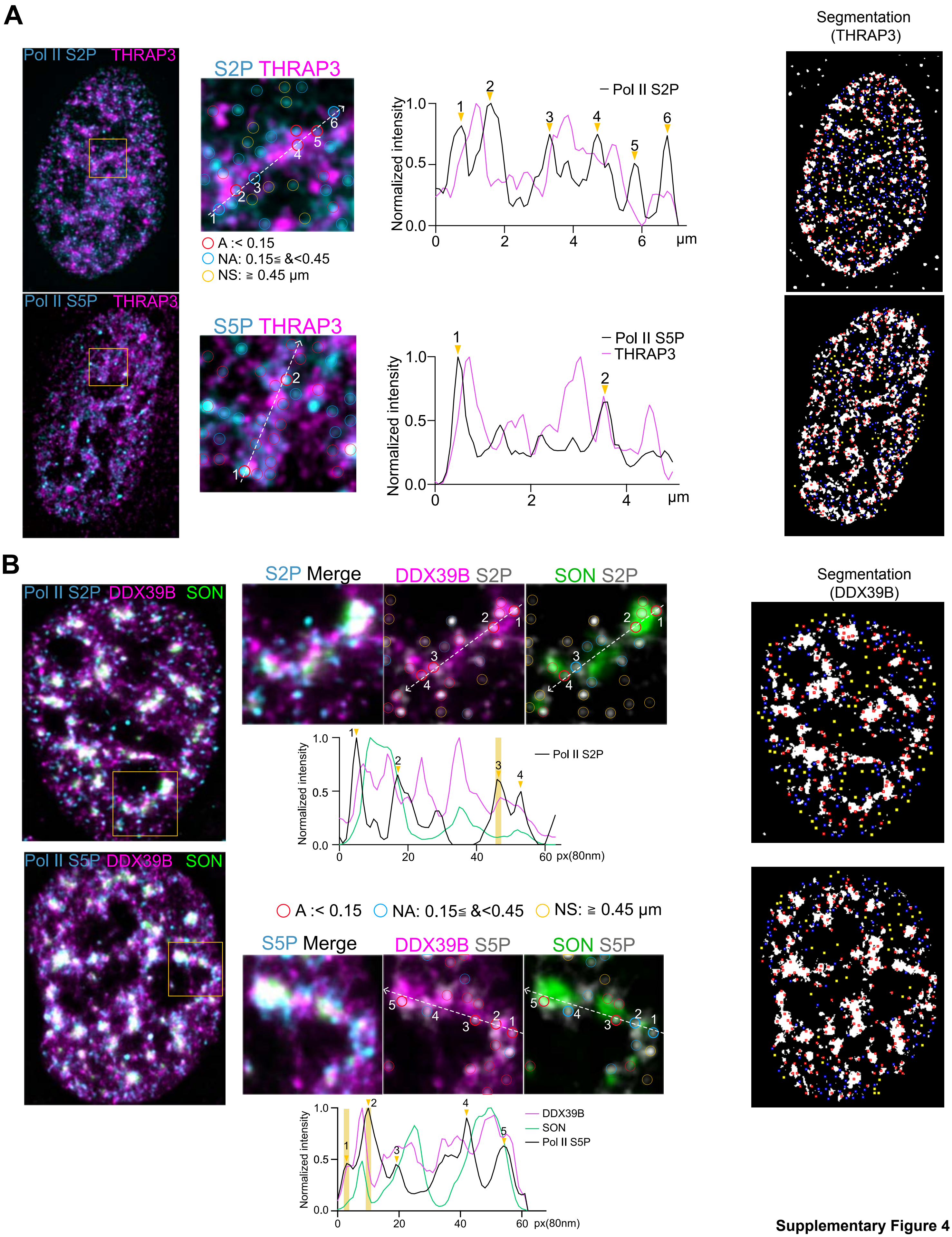


**Supplementary Figure 4**

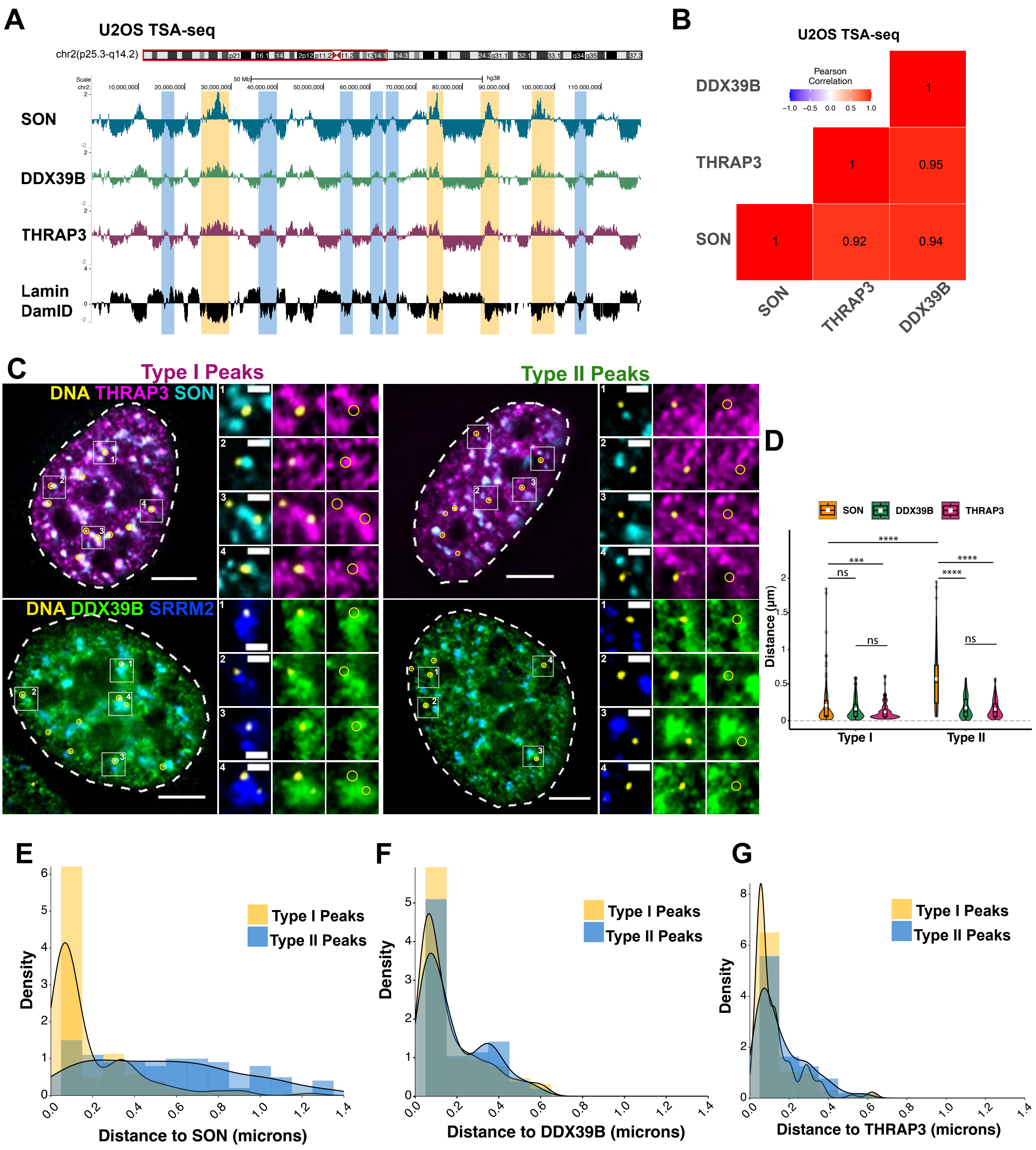


Suplementary Figure 5

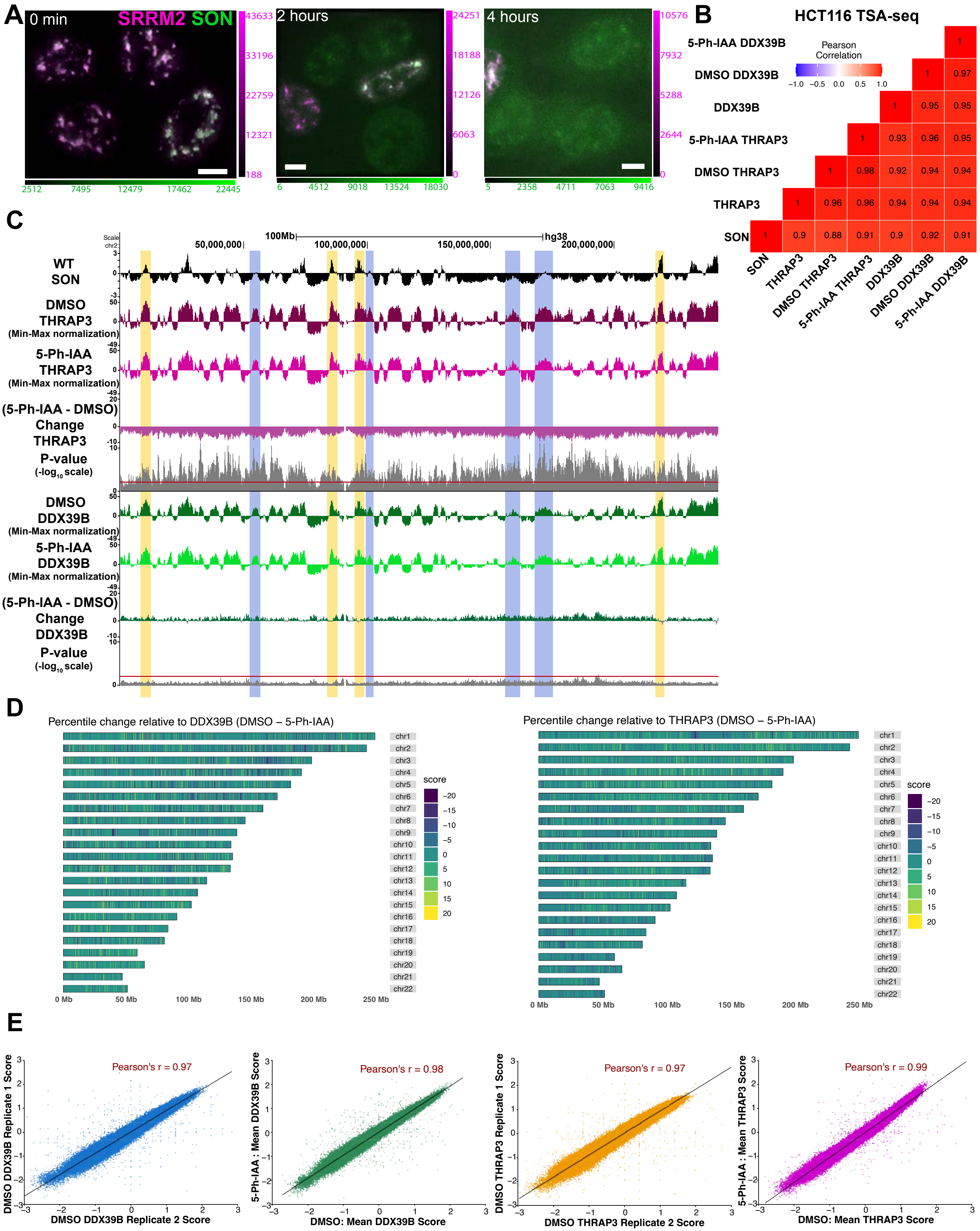


Supplementary Figure 8
